# Intercellular Mitochondrial Transfer as a Rescue Mechanism in Response to Protein Import Failure

**DOI:** 10.1101/2022.11.30.518494

**Authors:** Hope I Needs, Gonçalo C. Pereira, Emily Glover, Alina Witt, Wolfgang Hübner, Mark P. Dodding, Jeremy M Henley, Ian Collinson

**Author notes:** MRC - Mitochondrial Biology Unit, University of Cambridge, The Keith Peters Building, Cambridge Biomedical Campus, Cambridge CB2 0XY, UK.

## Abstract

Mitochondria are the powerhouses of eukaryotic cells, composed mostly of nuclear-encoded proteins imported from the cytosol. Thus, problems with the import machinery will disrupt their regenerative capacity and the cell’s energy (ATP) supplies–particularly troublesome for energy demanding cells like neurons and myocytes. Unsurprisingly then, dysfunctional import is implicated in disease. This study explores the consequences of import failure in mammalian cells; wherein, blocking the import machinery has profound effects on mitochondrial ultra-structure and dynamics, but, surprisingly, does not impact import. The explanation is an astonishing response involving intercellular mitochondrial transfer *via* tunnelling nanotubes: for the import of healthy mitochondria and jettisoning of those with jammed import sites. These observations support the existence of a widespread mechanism for the rescue of mitochondrial protein import failure.

**One-Sentence Summary:** A mitochondrial import rescue mechanism involving intercellular mitochondrial transport through tunneling nanotubes (TNTs).

## Main Text

Mitochondria are semi-autonomous organelles, found inside eukaryotic cells, responsible for oxidative phosphorylation and metabolic regulation amongst other functions. Their small genome encodes only a few proteins (13 in humans) while the remaining proteome (>1000 proteins) is synthesised in the cytosol and imported *via* highly specialised protein translocation machineries. The major route is the presequence pathway, taken by precursor proteins usually (∼70%) with cleavable N-terminal mitochondrial targeting sequences (MTS) [1]. These proteins are recognised at the surface of the outer mitochondrial membrane (OMM) by receptors of the translocase of the outer membrane (TOM) complex. Transport then proceeds through the TOM40 channel into the intermembrane space (IMS), towards the translocase of the inner membrane 23 (TIM23) complex [2, 3]. Additional internal sorting sequences are also deployed, which target the translocating precursor through the TIM23^SORT^ complex to be inserted laterally into the inner mitochondrial membrane (IMM) [2]. Otherwise, transport proceeds across the IMM and into the matrix *via* the TIM23^MOTOR^ complex. Passage of proteins into and across the IMM both require the membrane potential (ΔΨ), while pulling the protein into the matrix also requires ATP turnover [2]. MTS cleavage then enables release, folding, and assembly of the mature protein.

The consequences of mitochondrial import dysfunction, and thereby biogenic failure, are catastrophic for cell health; particularly for high energy consuming cells such as those of muscle and nervous tissue. Indeed, there are strong links between mitochondrial import failure and neurodegeneration, see [4] and references therein. Deficient import leads to the build-up of cytosolic precursor proteins, inducing the inhibition of protein synthesis, activation of the proteasome, and mitoprotein-induced stress responses [5–9]. While these responses have been shown to mitigate against the potential toxic effects of large quantities of aggregation prone precursor, they do not always successfully resolve the underlying problem of cells incapable of mitochondrial regeneration. To maintain overall bioenergetic function, mitochondria with failing import sites either need to be rectified or replaced. This paper describes a remarkable rescue mechanism in cells subject to perturbation of the mitochondrial import machinery. This response invokes the formation of tunneling nanotubes (TNTs), enabling intercellular mitochondrial transport to transfer viable mitochondria from heathy cells to replace defective mitochondria in compromised cells, as well as transferring compromised mitochondria into neighboring healthy cells.

There are multiple possible causes of mitochondrial import failure. Age, exposure to environmental and endogenous (*e.g.* reactive oxygen species–ROS) toxins, as well as nuclear and mitochondrial mutations can all impair bioenergetic function, resulting in depletion of ΔΨ and ATP required for import. Additionally, the TOM and TIM complexes can become blocked, *e.g.,* by stalled/misfolded precursor proteins. Indeed, refined kinetic analysis of protein translocation through the mitochondrial import and bacterial secretion machineries [3, 10, 11] show that transport failure and blockages are an integral component of the reaction cycle. Another possibility is that aggregation prone proteins, such as those variants implicated in neurodegenerative disease (amyloid precursor protein, alpha-synuclein, Huntingtin, and Tau) associate with the import machinery and perturb the transport process [12–14].

To study the consequences of import failure, and the response, we established tools to monitor and artificially block mitochondrial protein import in intact cells. HeLa cells were conditioned in galactose (HeLaGAL) to maximise their dependency on mitochondrial oxidative phosphorylation, in contrast to those grown in glucose (HeLaGLU), which favour glycolytic metabolism [15]. These cells were then utilised to study the impact of precursor stalling. Precursor proteins are imported in an unfolded state, so the C-terminal fusion of dihydrofolate reductase (DHFR), which resists unfolding when bound to the drug methotrexate (MTX), can be exploited to block the import machinery (Fig. 1A) [16, 17]. We therefore engineered a precursor construct encoding the MTS of the ATP synthase Subunit 9 (Su9) fused to a fluorescent reporter protein (EGFP or mScarlet) and DHFR for overexpression in HeLaGAL cells. Imaging and biochemical analysis confirmed that the mature protein was successfully targeted to mitochondria in the absence of MTX, but to a much lesser extent in its presence (Fig. 1A; S1 and S2). Interestingly, within the same cell, structured illumination microscopy (SIM) showed that some mitochondria (43%) were enveloped by aggregated precursor protein, while others were not (Fig. 1B–white and yellow arrows, respectively; S3 and S4).

**Fig. 1.**
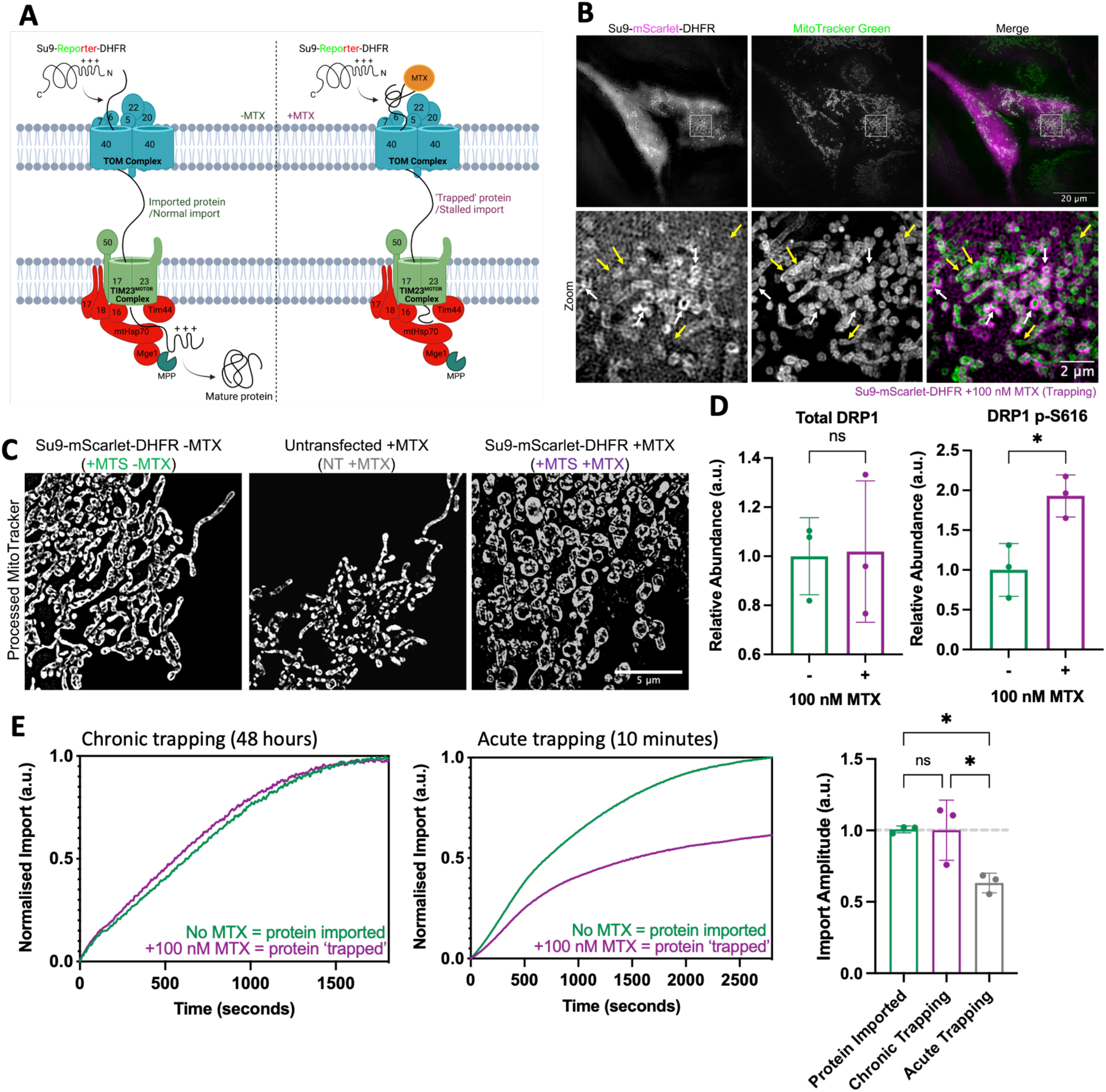
Chronic precursor trapping induces mitochondrial fission but has no impact on import. **(A)** In the absence of MTX (*left*), the precursor protein (*Su9-Reporter-DHFR*) is imported into mitochondria *via* the presequence pathway. In the mitochondrial matrix, the MTS is cleaved by MPP, and it folds to form the mature protein. In the presence of MTX (*right*), MTX binds to DHFR in the cytosol, preventing it from unfolding, and thus preventing it from crossing the TOM40 channel. Since the protein contains a long Su9-Reporter region before the DHFR, this region forms a plug through the TOM40 and TIM23 channels, mimicking a stalled or aggregated precursor protein within the presequence pathway. *Schematic created using BioRender*. **(B)** Representative SR 3D-SIM images showing trapping substrate accumulation around mitochondrial membranes in HeLaGAL cells expressing *Su9-mScarlet-DHFR* (magenta; 48h) in the presence of 100 nM MTX. Mitochondria are shown by MitoTracker Green staining (green). White arrows show association of trapping construct with mitochondria, yellow arrows show mitochondria not surrounded by precursor. Images are representative of N=3 biological replicates. **(C)** Representative SR 3D-SIM images showing mitochondrial morphology of HeLaGAL cells subjected to trapping (+MTS +MTX) or not (+MTS -MTS). Untransfected cells in the same well as cells subjected to trapping were used as an internal control for MTX treatment (NT +MTX). Images are representative of N=3 biological replicates. Unprocessed and uncropped images are shown in supplementary Fig. S5, quantified in S6. **(D)** Quantification of Western blots showing abundance of total DRP1 (*left*) or phospho-DRP1 S616 (*right*) in whole cell lysates from HeLaGAL cells expressing *Su9-EGFP-DHFR* in the absence or presence of 100 nM MTX. Normalised to β-actin loading control and -MTX control. N=3 biological replicates. Error bars represent SD. Representative blots are shown in supplementary Fig. S7. **(E)** NanoLuc import traces after precursor trapping. For chronic trapping (*left panel*), *Cox8a-11S* expressing HeLaGAL cells were subjected to expression of *Su9-mScarlet-DHFR* in the presence (purple; trapped precursor) or absence (green; imported precursor) of 100 nM MTX for 48 hours. For acute trapping (*middle panel*), HeLaGAL cells expressing *Cox8a-11S* were permeabilised using 3 nM rPFO prior to incubation (RT, 10 min) with 1 µM trap substrate (*Su9-ACP1-DHFR*), in the presence (purple; trapped precursor) or absence (green; imported precursor) of 100 nM MTX. NanoLuc import assays were then carried out to monitor import of precursor protein *Su9-EGFP-pep86*. Traces were normalised to eqFP670 fluorescence (as a marker of *Cox8a-11S* expression levels), technical replicates averaged, then normalised to the maximum value for the given run, and finally normalised to the +/-MTX (-MTS) control. Maximum import (amplitude) is shown (right panel). One-way ANOVA and Tukey’s *post-hoc* test were used to determine significance. Error bars represent SD. N=3 biological replicates, each with n=3 technical replicates.

High-resolution SIM revealed that artificial precursor trapping elicited a profound change in mitochondrial morphology (Fig. 1C; S5). The mitochondria became less branched, more rounded, and increased in width, indicative of fragmentation (Fig. 1C; S6). These effects did not occur when the precursor construct was not transfected, so they were not due to exposure of the drug MTX (Fig. 1C; S5 and S6; NT +MTX). We next analysed the abundance of a known regulator of mitochondrial fission, DRP1, which assembles into spiral-like oligomers when recruited to the OMM. Constriction brings about fission, mediated by GTP hydrolysis [18] and regulated by various post-translational modifications, including phosphorylation of DRP1 at serine616 (p-DRP1 S616) [19]. Western blotting showed that the abundance of total DRP1 was unchanged in HeLaGAL cells following mitochondrial precursor stalling, but phosphorylation at the pro-fission site S616 was increased (Fig. 1D; S7). Thus, we conclude that enhanced mitochondrial fission caused by precursor stalling accounts for the observed morphological changes.

We previously developed a split luciferase assay to monitor protein import into isolated yeast mitochondria [20], which we successfully adapted to our HeLa cell system [21]. Briefly, HeLa cells were cultured with the large fragment of the luciferase (11S) segregated in the mitochondrial matrix, so that the import of a precursor with the small fragment (pep86–required for luminescence) fused to the C-terminus can be monitored. This new capability enabled the reporting of the import of another precursor in HeLaGAL cells subject to precursor stalling by DHFR-MTX, after plasma membrane permeabilisation.

We first tested import dynamics after chronic exposure to precursor stalling (48 hours of *su9-mScarlet-DHFR* overexpression +MTX; see above). Surprisingly, we observed no effect on the import of a second precursor protein *Su9-EGFP-pep86* in our in-cell (HeLaGAL) assay (Fig. 1E–left). However, under more acute stalling conditions *via* the addition of high concentrations of purified precursor protein-DHFR fusion in the presence of MTX and NADPH (for increased stabilisation of DHFR) for 10 minutes, the import of the reporter precursor was diminished (Fig. 1E–middle). Since analogous treatments of isolated mitochondria *in vitro* completely blocked the import sites [3], the preservation of mitochondrial import inside cells following chronic precursor trapping was unexpected. Based on these findings we hypothesised that a clearance or rescue mechanism must be operating, but lost following mitochondrial isolation.

We observed that the cells subject to chronic exposure to precursors with DHFR and MTX (chronic precursor import stalling), exhibited a dramatic increase in thin elongated protrusions (Fig. 2A; S8). The increase in protrusions was dependent on DHFR-induced precursor stalling, and not an off-target effect of MTX, because they were not apparent when the MTS was omitted. Furthermore, this occurrence was dependent on galactose conditioning, confirming the response is mediated by mitochondria [15] (Fig. 2B; S8). Immunocytochemistry and confocal microscopy showed that the protrusions contain microtubules and actin (Fig. 2C), characteristic of tunnelling nanotubes (TNTs), capable of organellar (including mitochondrial) transport between cells [22–25]. To further investigate the identity of these protrusions, cells were treated with nocodazole, which impairs TNT establishment through inhibition of microtubule formation [26–30]. In doing so, the number of protrusions was indeed reduced to the baseline levels (Fig. 2D; S9), consistent with their identity as such.

**Fig. 2.**
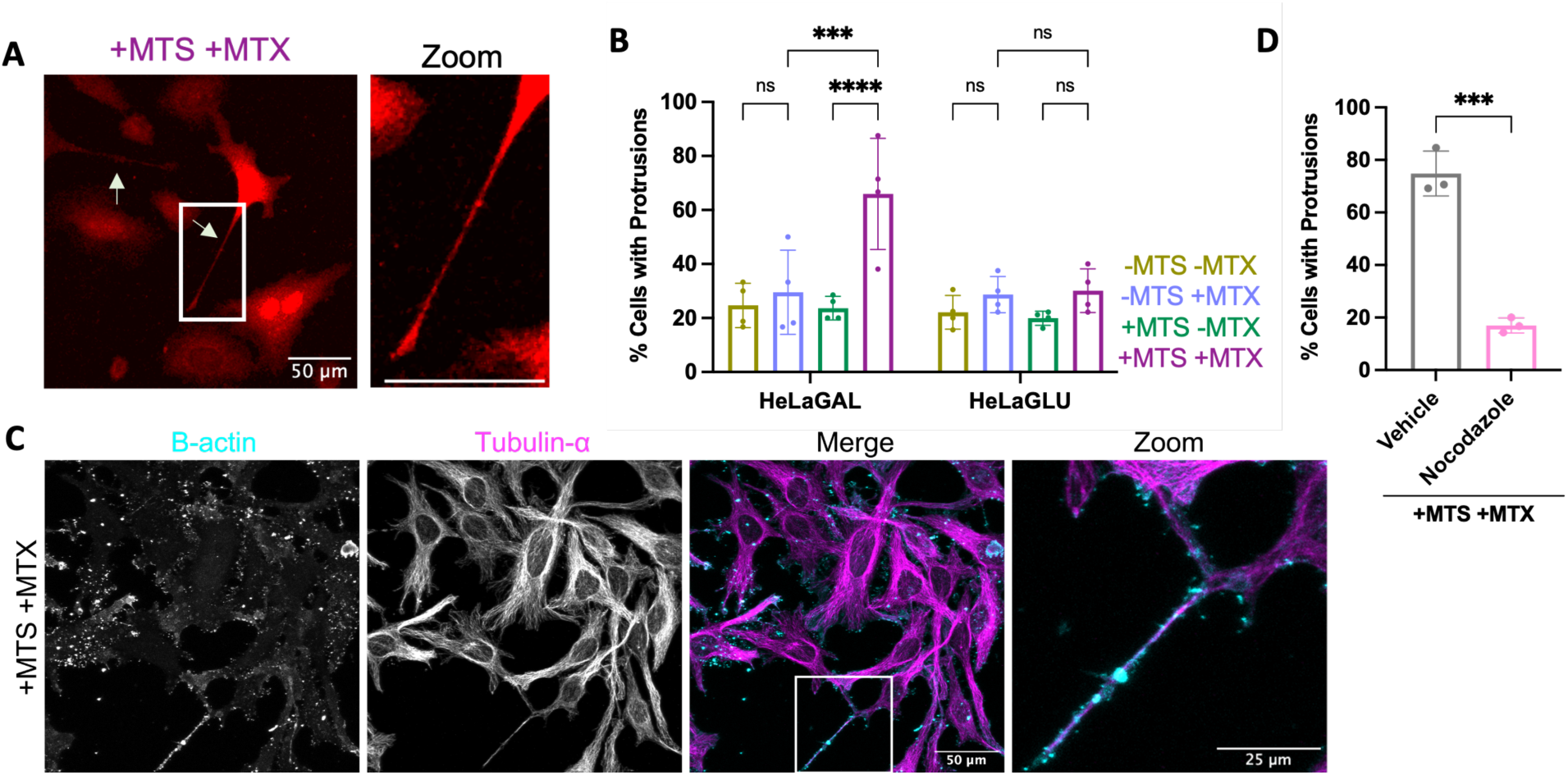
Chronic precursor trapping induces formation of TNTs. **(A)** Representative confocal images showing cell morphology of HeLaGAL cells subjected to expression of *mCherry* (red) and *Su9-EGFP-DHFR* in the presence of 100 nM MTX (+MTS +MTX; trapped precursor). Arrows indicate protrusions, and box highlights zoom region. N=4 biological replicates. **(B)** Quantification of the proportion of transfected cells with protrusions. All representative images shown in supplementary Fig. S8. Error bars show SD. Two-way ANOVA and Tukey’s *post-hoc* test were used to determine significance. N=4 biological replicates; 20 cells counted per replicate. **(C)** Representative confocal images showing tubulin and actin staining of protrusions. HeLaGAL cells were subjected to trapping for 48 h to induce protrusion formation. Cells were fixed and stained with antibodies against β-actin (Sigma; A2228) and tubulin-α (BioRad; MCA78G) and visualised by confocal microscopy to observe protrusion composition. N=4 biological replicates. **(D)** Quantification of the proportion of HeLaGAL cells, subjected to precursor trapping, with protrusions following treatment for 48 hours in the absence (vehicle; DMSO only) or presence of 100 nM nocodazole. Cells were pre-treated with 100 nM MTX for 16 h, then treated with nocodazole at the same time as transduction with the trapping precursor and incubated for 48 h prior to imaging. Representative confocal images shown in supplementary Fig. S9. Error bars show SD. Unpaired t-test was used to determine difference between groups. N=3 biological replicates; 20 cells counted per replicate.

We next investigated whether these TNTs could be capable of intercellular mitochondrial transport to recruit functional mitochondria, or remove those that have become compromised. Close inspection by super-resolution imaging highlighted the presence of mitochondria inside tubes connecting a cell challenged by precursor stalling with an untransfected healthy cell (Fig. 3A). To confirm mitochondria were transferred between cells, co-cultures of challenged HeLaGAL cells (producing a stalling precursor and a green mitochondrial marker: *Su9-EGFP*) and healthy cells (producing a red mitochondrial marker protein: *Cox8a-DsRed*) were established in the presence of MTX, and imaged by confocal microscopy. The results clearly identified cells containing a mixture of mitochondria originating from both healthy and challenged cells (Fig. 3B). Furthermore, live cell light-sheet fluorescence microscopy showed healthy mitochondria (without precursor stalling) being transferred through a TNT towards a challenged cell (Fig. 3C; supplementary Movie S1). This observation explains why, within a cell exposed to precursor trapping, some mitochondria are surrounded by aggregated precursors, and thereby stalled, while others in the same cell are not (Fig. 1B; S4), the latter potentially originating from TNT connected healthy cells.

**Fig. 3.**
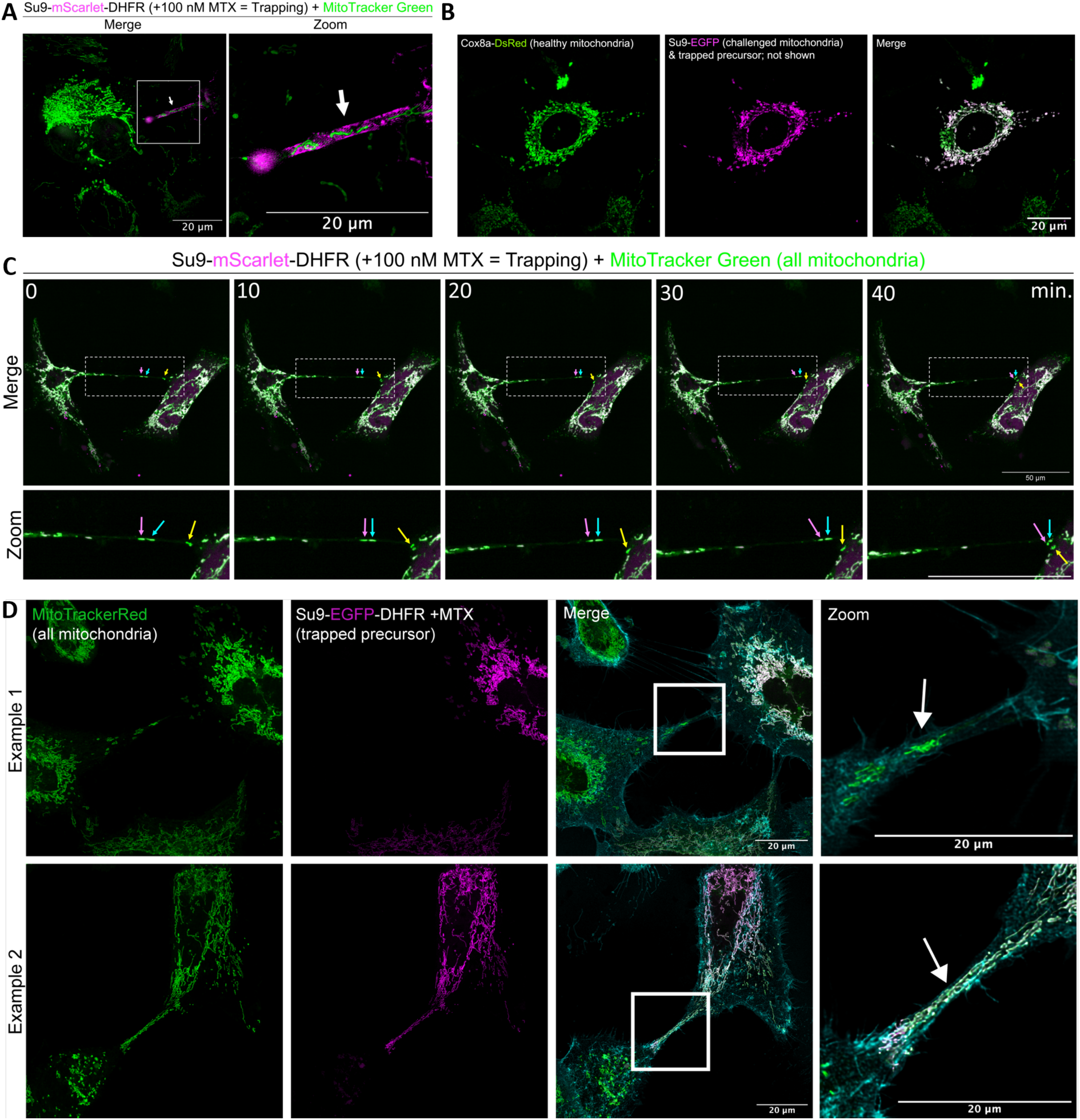
Chronic precursor trapping induces intercellular mitochondrial transfer *via* TNTs. **(A)** Representative SR 3D-SIM image showing mitochondria (MitoTracker Green; green) within a TNT formed between a HeLaGAL cell with trapped precursor (*Su9-mScarlet-DHFR*; magenta), and an untransfected neighbour (green mitochondria, unlabeled cytosol). Cells were pre-treated for 16 h with 100 nM MTX, transfected with *Su9-mScarlet-DHFR* and incubated for 48 hours prior to mitochondrial staining with MitoTracker Green and live imaging by SIM. Arrow highlights TNT and box indicates zoom area. N=3 biological replicates. **(B)** Representative confocal images from co-culture experiments, showing mitochondrial transfer between healthy and sick cells. Mitochondria of one batch of HeLaGAL cells were subjected to *Cox8a-dsRed* overexpression (healthy mitochondria; green). A second batch of cells was subjected to *Su9-EGFP* expression (magenta) and sequential trapping (*Su9-eqFP670-DHFR* +100 nM MTX). After 48 hours, cells were washed, detached, counted, and mixed at equal ratios, prior to reseeding on coverslips. Cells were co-cultured for a further 8 hours before fixation and visualisation by confocal microscopy. N=3 biological replicates. **(C)** Time series of still images from a representative time-lapse movie showing the transfer of healthy mitochondria (MitoTracker Green; green) within a TNT into a cell with trapped precursor (*Su9-mScarlet-DHFR* (magenta) +100 nM MTX). Cells were imaged live using an Olympus IXplore SpinSR system. Each timeframe is a Max z-projection of 10 slices, taken every 5 min x 10 timepoints. Full movie shown in supplementary Movie S1. **(D)** Representative confocal images showing mitochondria (MitoTracker Red; green) within a TNT (plasma membrane stained with Wheat Germ Agglutinin (WGA); cyan) formed between a HeLaGAL cell with trapped precursor (*Su9-EGFP-DHFR*; magenta), and an untransfected healthy neighbour (green mitochondria). Example 1 shows healthy mitochondria being transferred to an unhealthy cell; example 2 shows unhealthy mitochondria being transferred to a healthy cell. Cells were transfected with *Su9-EGFP-DHFR* and incubated for 48 hours in the presence of 100 nM MTX prior to staining with MitoTracker Red and WGA. Cells were then fixed and imaged. Box indicates zoom area and arrow highlights TNT. N=3 biological replicates.

Further analysis by confocal microscopy identified TNTs containing healthy mitochondria, travelling from a healthy cell to a cell with trapped precursor (Fig. 3D; example 1). Conversely though, there were other instances where TNTs were observed containing mitochondria originating from cells with trapped precursor (Fig. 3D; example 2). Thus, it appears that mitochondria travel in both directions; potentially for replenishment from healthy cells, and for affected cells to jettison impaired mitochondria. Interestingly, while mitochondria with trapped precursors are subject to TNT transport, cytosolic precursor proteins accumulated in the cytosol are not – suggesting transport occurs actively, rather than passively by diffusion. To explore whether this process is a general response to perturbation of the protein import apparatus, HeLa cells were exposed to MitoBloCK-20 (MB20) – a small molecule inhibitor of the presequence pathway acting on TIM17, a key component of the TIM23 complex [31]. Similar to precursor stalling, only acute exposure to the drug affected protein import activity (Fig. 4A). The failure of chronic drug exposure to impact protein import coincides with a large increase in the proportion of cells with structures resembling TNTs (Fig. 4B; S10); once again this only occurred in cells dependent on mitochondria for ATP synthesis (HeLaGAL). These TNTs, induced by chemical inhibition, also contained actin, microtubules, and mitochondria (Fig. S11).

**Fig. 4.**
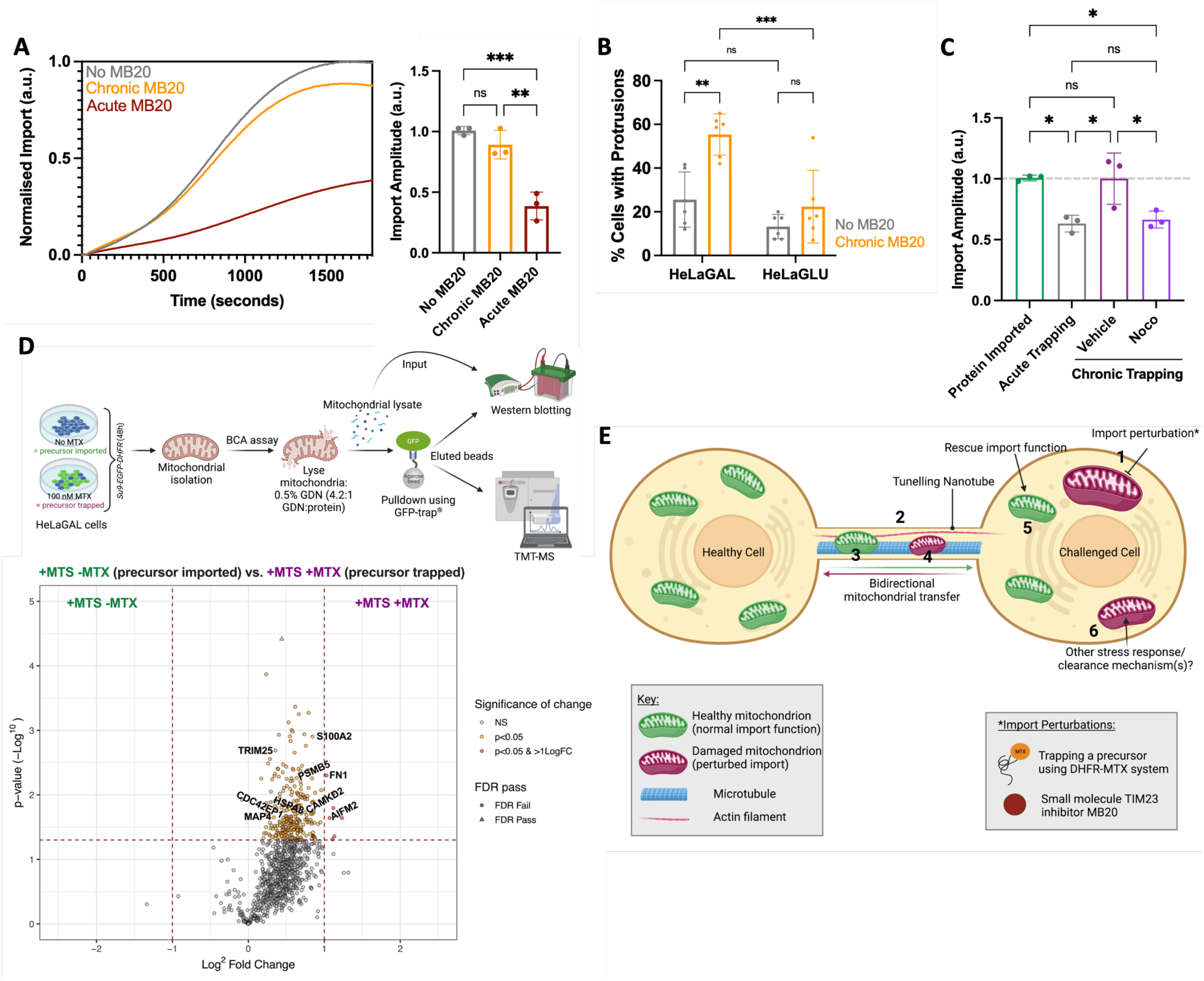
Intercellular mitochondrial transfer *via* TNTs rescues import function. **(A)** NanoLuc import traces showing precursor protein (*Su9-EGFP-pep86*) import in *Cox8a-11S* expressing HeLaGAL cells treated with vehicle only (DMSO; no MB20; grey), or with either chronic (48 h; orange) or acute (10 min; red) MB20 exposure. Quantification of import amplitude is shown in right hand panel. N=3 biological replicates, each with 3 technical replicates. Error bars represent SD. One-way ANOVA with Tukey’s *post-hoc* test was used to determine significance. **(B)** Quantification of proportion of cells with TNTs following chronic treatment with 10 µM MB20 (48 h). Representative confocal images shown in supplementary Fig. S10. N=6 biological replicates; 20 cells counted per replicate. Error bars represent SD. Two-way ANOVA and Tukey’s *post-hoc* test were used to determine significance. **(C)** Quantification of import amplitude in cells subjected to trapping, in response to treatment with nocodazole. HeLaGAL cells expressing *Cox8a-11S* were subjected to trapping (+MTX) or not (-MTX) and treated with 100 nM nocodazole (*noco*), or DMSO only (*vehicle*) for 48 hours prior to monitoring the import of *Su9-EGFP-pep86* by NanoLuc import assays. Import trace is shown in supplementary Fig. S12. N=3 biological replicates, each with 3 technical replicates. Error bars represent SD. One-way ANOVA with Tukey’s *post-hoc* test was used to determine significance. **(D)** HeLaGAL cells were subjected to *Su9-EGFP-DHFR* expression in the presence or absence of 100 nM MTX for 48 h. Mitochondria were isolated and mitochondrial protein content was assessed using a BCA assay. Equal amounts of mitochondria were lysed using 0.5% GDN at a ratio of 4.2:1 (GDN: protein). A fraction of mitochondrial lysate was saved as an input control, and the rest was loaded onto GFP-trap agarose beads, to pull-down proteins associated with the GFP tagged protein of interest (trapped or imported precursor). The resulting beads conjugated to proteins associated with GFP-tagged proteins were analysed by Western blot and TMT-MS. Lower panel shows a volcano plot highlighting proteins enhanced in pulldown samples from mitochondria with imported precursor (+MTS -MTX, *left*) compared to those with the trapped precursor (+MTS +MTX, *right*). Selected proteins of interest are highlighted by gene names on the volcano plot and the full list is provided in supplementary Table S1. A representative Western blot is shown in supplementary Fig. S13 and control (-MTS, +/-MTX) volcano plot in supplementary Fig. S14. N=3 biological replicates. *Top panel schematic created using BioRender*. **(E)** Proposed import rescue mechanism *via* TNT-dependent intercellular mitochondrial transfer. Cells subjected to import perturbations *(1)* induce the formation of TNTs composed of microtubules and actin filaments *(2).* TNTs allow transfer of healthy mitochondria from neighboring cells into cells with challenged mitochondria *(3),* whilst also allowing clearance of damaged mitochondria to neighboring healthy cells *(4)*. The transfer and integration of healthy mitochondria with normal import function, and disposal of mitochondria with defective import, rescues the overall cellular import function *(5)*. This rescue effect is likely enhanced by additional stress response mechanisms *(6). Schematic created using BioRender*.

The results presented so far suggest that the TNTs are part of a widespread rescue mechanism that enables cells containing mitochondria with compromised import to receive functional mitochondria from nearby healthy cells, and unload dysfunctional mitochondria for degradation/regeneration. To test this hypothesis, we re-examined protein import, which was apparently unaffected in cells subjected to chronic precursor stalling (Fig. 1E), but in the presence of TNT inhibitor nocodazole. In cells exposed to chronic precursor trapping and treated with nocodazole, we observed a reduction in the total amount of protein imported compared to cells without trapped precursor (Fig. 4C; S12). Therefore, the unexpected resilience of import function, within cells subject to precursor trapping, is dependent on TNTs.

To investigate this rescue mechanism further we deployed tandem mass tagging mass spectrometry (TMT-MS) to identify proteins associated with the trapped precursor. Cells were subject to stalling again through production of a *Su9-EGFP-DHFR* fusion (with and without the MTS and MTX as controls). Following a chronic (48 hour) treatment, mitochondria from HeLaGAL cells were isolated and proteins gently extracted with glyco-diosgenin amphiphile (GDN). Proteins interacting with the precursor were then isolated using a GFP-trap, which effectively binds the precursor (Fig. 4D; S13). TMT-MS analysis of the captured precursor and its associates shows a significant enhancement of proteins when derived from cells cultured in the presence, compared to those in the absence, of MTX. The volcano plot (Fig. 4D; supplementary Table S1) highlights proteins in complex with the stalled precursor (+MTX), controlled against those associated with the imported form (-MTX).

Of the significant hits with >2 fold enhancement, cell adhesion protein fibronectin (FN1) has been shown to be critical for TNT formation and activity [32], whilst apoptosis inducing factor mitochondria associated 2 (AIFM2) is related to mitochondrial import stress signalling [33], and is induced by p53 activity [34] which can induce TNT formation [35]. Interestingly, there is also an enhancement of calcium signalling proteins Calcium/calmodulin-dependent protein kinase type II subunit delta (CamKIID) and S100A2. Calcium signalling *via* the Ca^2+^/Wnt pathway has been shown to regulate TNTs previously [36]. Additionally, cytoskeleton related proteins required for TNT formation, such as Microtubule associated protein 4 (MAP4) and Cdc42 effector protein 1 (CDC42EP1), were enhanced [37–39].

Additionally, associated proteins include several proteins involved in stress response pathways, particularly associated with the proteasome (*e.g.,* E3 ubiquitin ligase TRIM25 [40], Proteasome subunit β type 5 (PSMB5) [41], and Heat shock cognate 71 kDa protein (HSPA8) [42, 43]), suggesting that other mitoprotein-induced stress response pathways are also at play. An experiment was also conducted without an MTS to show the non-specific effects of MTX (Fig. S14). In this control there was only one significant enhancement in the presence of MTX (not present in +MTS +MTX); therefore, the hits identified as significantly enhanced represent true associations with the trapped precursor.

Taken together, our results demonstrate that impaired mitochondrial import, be it precursor stalling at the outer membrane or inhibition of the TIM23 complex, invokes a rescue mechanism involving the intercellular transport of mitochondria *via* filamentous, membranous protrusions: TNTs that contain F-actin and microtubules. TNTs have been described previously for their involvement in long-range intercellular communication and transport; including of lipids, nucleic acids, microRNAs, ions such as calcium, fragmented plasma membrane, viruses (*e.g.* coronaviruses) [44] and even entire organelles: mitochondria, ER, Golgi, and endosomes [22-25, 45, 46]. The dimensions of TNTs seem to be tailored according to the cargo, with distinct thin and thick varieties, the latter with a diameter of 1-7 µm, such as those seen here for mitochondrial transport [47–49]. TNT formation can be induced by various stress factors including H_2_O_2_, serum depletion, mitochondrial stress, and cytotoxicity, which may act *via* the p53 pathway [35]; they alleviate cellular stress by enabling replenishment and/or removing harmful debris. Indeed, healthy mitochondria have previously been reported to transport through TNTs and rescue cells in the early stages of apoptosis [39]. Another study demonstrated that TNTs can assist in reducing cellular toxicity caused by alpha-synuclein fibrils (implicated in Parkinson’s disease) in microglia, by facilitating the removal of protein aggregates and the delivery of healthy mitochondria [50]. Here, we show how direct mitochondrial import impairment leads to bidirectional mitochondrial transfer *via* TNTs, suggestive of a widespread mechanism for rescue of mitochondrial function.

It makes sense that the TNT response described, triggered by failing import, helps resolve potentially catastrophic incapacitated mitochondrial biogenesis. Beyond that, we also speculate that the TOM and TIM23 complexes act as reporters of mitochondrial health; whereby, for whatever reason, reduced bioenergetic function would lead to diminished Δψ and/ or ATP, decreasing import efficiency and hence initiating rescue *via* TNTs. In this respect, the connection between protein aggregation associated with neurodegeneration and mitochondrial import is particularly intriguing. Indeed, the intercellular transport of the Tau protein and β-amyloid, associated with Alzheimer’s disease, can also occur *via* TNTs [51]. This could be connected to the propensity of these aggregation prone proteins to associate with the mitochondrial import machinery. Variants of the Huntingtin protein, the causative agent of disease, and the β-amyloid precursor APP have both been shown to associate with the TOM40/TIM23 complexes [12, 14, 52]. Indeed, in a parallel study we show that the a Tau variant linked with neurodegeneration associates with TOM40 and brings about TNT formation [53]. Therefore, these proteins may also act through binding to and impairing the import machinery, inducing TNT formation for aggregate efflux [54–56]. On a more sinister note, perhaps this mechanism could be hijacked to spread infectious agents; whereby viruses or prions target themselves to the mitochondrial import machinery and ride in, through TNTs, to other cells [44, 57, 58].

What emerges from our study is a picture of an astonishing process for rescue of mitochondrial import function involving intercellular mitochondrial transport through TNTs (Fig. 4E). It may well be one of many rescue mechanisms instigated by perturbation of the import apparatus, including the stress response to unfolded precursors in the cytosol [5, 6]. It will be very interesting to see how this mechanism is manifested *in vivo*, to what extent and for what purpose, since mitochondrial import dysfunction underpins a wide range of devastating and often fatal diseases [4, 59, 60]. Greater understanding of the formation and functions of TNTs will provide exciting opportunities for biological engineering and for medical applications. For example, modulating this process genetically, or with small molecules, as we have demonstrated, could enable targeted mitochondrial transport *in vivo*, and approaches to mitochondrial replacement therapy for cell and tissue regeneration.

## Acknowledgments

We gratefully acknowledge access and support of the Wolfson Bioimaging Facility for imaging support. We are grateful to Dr Stephen Cross and Dr Richard Seager for image analysis support. We gratefully acknowledge the University of Bristol Proteomics Facility, specifically Dr Kate Heesom, for support with TMT-MS, and Dr Phil Lewis for bioinformatics support.

## Funding

This work was funded by the Welcome Trust: through the Wellcome Trust *Dynamic Molecular Cell Biology* PhD programme (HIN, 083474; EG, 218510) and a Wellcome Investigator award (IC and GCP, 104632).

The funders had no role in study design, data collection and interpretation, or the decision to submit the work for publication. For the purpose of Open Access, the authors have applied a CC BY public copyright license to any Author Accepted Manuscript version arising from this submission.

## Author contributions

HIN, EG, GCP, MPD, JMH and IC designed experiments; HIN, EG, AW and WH conducted experiments; HIN, GCP, JMH and IC wrote the manuscript; IC and JMH secured funding and led the project.

## Competing interests

The authors declare no competing interests.

## Data and materials availability

All data are available in the main text or the supplementary materials.

## Supplementary Materials

### Materials and Methods

#### Reagents

All chemicals were of the highest grade of purity commercially available and purchased from Sigma, UK, unless stated otherwise. Aqueous solutions were prepared in ultrapure water, while for non-aqueous solutions, ethanol or DMSO was used as solvent instead. Antibody catalogue numbers and suppliers are detailed in figure legends.

#### Generation of Constructs

Constructs were generated by standard cloning techniques. Briefly, PCR reactions were carried out using Q5 High Fidelity Hot Start DNA Polymerase (New England Biolabs (NEB)), using 20 pmol primers and 200 pg template DNA, as per manufacturers’ instructions. PCR products were purified using QIAquick PCR Purification Kit (QIAgen). Restriction digest reactions were carried out using NEB restriction enzymes at 37°C for 45 min. Ligation reactions were carried out using T4 DNA Ligase (NEB) overnight at 16°C. Transformation was carried out in *E. coli* cells (α-select, XL1-Blue, or BL21-DE3 cells were used, depending on application; all originally sourced from NEB) for 30 min on ice, followed by heat shocking (45 sec, 42°C), and a further 15 min on ice. Cells were recovered by incubating in LB media at 37°C for 1 h and then plated on LB-agar plates containing appropriate antibiotic. Plasmids were prepared by mini or maxi preps using commercially available kits (Qiagen and Promega, respectively) following manufacturers’ instructions, and verified by DNA sequencing using Eurofins Genomics TubeSeq service.

#### Protein Purification

Protein expression was carried out exactly as described previously [20]. A single colony of transformed BL21 (DE3) bacteria was grown as a pre-culture in LB with appropriate antibiotic overnight (37°C; 200 rpm). Pre-cultures were used to inoculate a secondary culture at a 1:100 dilution, in 2X YT supplemented with appropriate antibiotic. Secondary cultures were grown until mid-log phase, then induced with 1 mM IPTG or 0.2% (w/v) Arabinose and grown for a further 3 h. Cells were harvested by centrifugation (15 min; 6000 xg), resuspended in TK buffer (20 mM TRIS base, 50 mM KCl; pH 8.0), cracked in a cell disruptor (Constant Systems; 2 cycles at 25 kpsi), and clarified by centrifugation (45 min; 38,000 rpm).

##### GST tagged Recombinant Perfringolysin (rPFO)

Supernatant (soluble fraction) was loaded onto a 5 mL GSTrap 4B column (GE Healthcare) and the column was washed with TK buffer until the absorbance of the flow through ceased to decrease any further. The peptide was eluted using 10 µM reduced glutathione, prepared fresh. Eluted fractions were pooled and loaded onto a 5 mL anionic exchanger (Q-column; GE Healthcare). A salt gradient of 0-1 M was applied over 20 min and the protein was eluted in 5 mL fractions. The fractions containing the protein were confirmed by SDS-PAGE with Coomassie staining, then pooled and spin concentrated. The final protein concentration was determined based on an extinction coefficient of 117,120 M^-1^ cm^-1^. The protein was aliquoted, snap frozen, and stored at −80°C until required. For each assay, a fresh aliquot was thawed, used immediately, and discarded afterwards, due to the instability of this protein [61].

##### GST-Dark Peptide

GST-Dark was prepared as described previously [20]. Supernatant was loaded onto a GSTrap 4B column and purification was carried out exactly as for rPFO but without the ion exchange chromatography. Analysis, yield, and freezing was carried out exactly as for rPFO. Protein concentration was determined based on an extinction coefficient of 48,360 M^-1^ cm^-1^.

##### His tagged Su9-EGFP-pep86

Inclusion bodies (insoluble fraction) were solubilised in TK buffer supplemented with 6 M urea, before loading onto a 5 mL HisTrap HP column (GE Healthcare). The protein was eluted in 300 mM imidazole. Eluted fractions containing the desired protein were pooled and loaded onto a 5 mL cationic exchanger (S-column; GE Healthcare). A salt gradient of 0-1 M was applied over 20 min and the protein was eluted in 5 mL fractions. Imidazole was removed by spin concentration, followed by dilution in TK buffer containing 6 M urea. Analysis, yield, and freezing were carried out exactly as for rPFO. Protein concentration was determined based on an extinction coefficient of 28,880 M^-1^ cm^-1^.

##### His-SUMO--Su9-ACP1-D-ACP1-D-DHFR-Myc

His-SUMO--Su9-ACP1-D-ACP1-D-DHFR-Myc was produced as described previously [3]. It was expressed by induction with IPTG, and cells were grown for a further 18 hours prior to harvesting. Subsequently, proteins from the soluble fraction were purified using a Nickel column, and finally contaminants were removed by size exclusion chromatography using a HiLoad 16/60 Superdex gel filtration column (Cytiva, UK). The SUMO tag was cleaved from the protein using Ulp1 protease following purification, and the eluted sample passed through a Nickel column to separate the SUMO-His tag from the protein. Protein concentration was determined based on an extinction coefficient of 42,400 M^-1^ cm^-1^.

#### Cell Culture

HEK293T (ECACC) and glucose-grown HeLa cells (HeLaGLU; ATCC) were maintained in Dulbecco’s Modified Eagle’s Medium (DMEM; Gibco; 41965039) supplemented with 10% (v/v) foetal bovine calf serum (FBS; Invitrogen) and 1% (v/v) penicillin-streptomycin (P/S; Invitrogen). When OXPHOS dependence was required, HeLa cells were cultured in galactose medium (HeLaGAL; ATCC) consisting of DMEM without glucose (Gibco; 11966025) supplemented with 10 mM galactose, 1 mM sodium pyruvate, 10% FBS and 1% P/S. Cells were cultured in galactose media for at least 3 weeks prior to experiments on HeLaGAL cells. Cells were maintained in T75 ventilated flasks in humidified incubators at 37°C with 5% CO_2_.

#### Cell Transduction by Transfection

HeLa cells were plated and grown up to ∼70-80% confluency. At this point cells were transfected with 1 μg (per 35mm dish) of the desired DNA using Lipofectamine 3000 reagent (Thermo Fisher Scientific), at a 1:1.5 ratio of DNA:Lipofectamine, following the manufacturers’ protocol. Cells were then grown for a further 24-72 h prior to experimental analysis.

#### Cell Transduction by Lentiviral Infection

Lentiviral particles were produced in HEK293T cells by addition of a mixture of DNAs (27.2 µg DNA to be produced, and packaging vectors pMDG2 (6.8 µg) and pAX2 (20.4 µg)) and 1 µg/µL pEI transfection reagent in OptiMEM media (Gibco) to HEK293T cells in a T75 flask, followed by incubation for 6 hours at 37°C, 5% CO_2_. Media was then changed to complete DMEM, and cells incubated for 72 h to allow lentivirus particle production. Lentiviral particles were harvested at 48 and 72 h for maximum yield, pooled, spun down at 4000 xg for 5 min to remove dead cells, and concentrated by adding Lenti-X concentrator (Takara Bio) at a 1:3 ratio and incubating at 4°C for at least 1 h. Lentivirus was then pelleted by centrifugation at 4000 xg for 45 min. Pellets were resuspended in plain DMEM at 1:50 of initial supernatant volume and aliquoted and stored at −80°C until required. For infection, concentrated lentivirus (volume optimised by titration of each fresh batch) was added dropwise to cell media when cells were at ∼70-80% confluency and incubated for 24-72 h prior to experimental analysis.

#### Total Protein Cell Lysis

For extraction of total protein lysate, cells were washed extensively with HBSS, and RIPA buffer (Sigma, supplemented with 1 mM PMSF) was added (200 µL for ∼35 mm surface, scaled up or down appropriately). Cells were scraped on ice into Eppendorf tubes, which were incubated for 1 h on a rotating wheel at 4°C, prior to spinning down at 10,000 xg for 15 min at 4°C. Supernatant (containing proteins) was stored at - 20°C. If protein concentration was required, this was obtained by carrying out a BCA assay (Pierce^TM^ BCA Protein Assay Kit, Thermo Fisher Scientific), following manufacturer’s instructions and using BSA as a standard.

#### Western Blotting

Following total protein extraction, protein samples were heated to 95°C for 5 min in the presence of LDS supplemented with 50 mM DTT. 30 μg of total protein was loaded on 4-12% BOLT gels (Thermo Fisher Scientific), separated (200 V, 24 min), and transferred onto polyvinylidene difluoride (PVDF) membranes (activated with methanol; Thermo Fisher Scientific) with transfer buffer (336 mM tris, 260 mM glycine, 140 mM tricine, 2.5 mM EDTA) using a semi-dry Pierce Power Station transfer system (Thermo Fisher Scientific; 25 V, 2.5 mAmp, 10 min). Membranes were blocked for 1 h in milk or BSA (5% w/v in TBS-T: 20 mM TRIS, 1.5 M NaCl, 0.1% (v/v) Tween-20 (pH 7.6)) and incubated in 5% milk or BSA containing the appropriate primary antibody (see figure legends; 4°C, overnight). Membranes were washed extensively with TBS-T and probed with appropriate secondary antibody in 2.5% milk or BSA (RT, 1 hour). Membranes were washed with TBS-T, incubated with ECL substrate (GE Healthcare), and developed using an Odyssey Fc Imaging System (LI-COR). Analysis and quantification were carried out using Image Studio Lite software

#### Mitochondrial Isolation

Confluent cells were harvested by trypsinisation, pelleted, and washed extensively with HBSS. Pellets were frozen overnight at −80°C and thawed the next day, to weaken membranes. Subsequently, mitochondrial isolation was performed using Mitochondrial Isolation Kit for Cultured Cells (Abcam; ab110170) following manufacturers’ instructions. Mitochondrial protein concentration was calculated by BCA assay.

#### Immunoprecipitation

Following mitochondrial isolation, mitochondria were gently lysed using 4.2g/g GDN (4.2g GDN: 1g protein) in IP buffer (0.1 M TRIS-HCl, 0.15 M NaCl, phospholipids (0.03mg/ml PE; 0.03mg/ml PG; 0.09mg/ml PC), 1X cOmplete ULTRA Protease Inhibitor Cocktail). GFP-tagged proteins were isolated using 10 µL GFP-trap beads (Chromotek). Supernatant (lysed mitochondrial sample) was incubated on a rotating wheel with beads overnight at 4°C. Subsequently, beads were washed in IP buffer. After washing, supernatant was removed, and samples were analysed by Western blotting or mass spectrometry.

#### Proteomic analysis

##### TMT Labelling and High pH reversed-phase chromatography

Immuno-isolated samples were reduced (10 mM TCEP, 55°C for 1 h), alkylated (18.75 mM iodoacetamide, room temperature for 30 min) and then digested from the beads with trypsin (2.5 µg trypsin; 37°C, overnight). The resulting peptides were then labeled with TMTpro sixteen-plex reagents according to the manufacturer’s protocol (Thermo) and the labelled samples pooled and desalted using a SepPak cartridge, according to the manufacturer’s instructions (Waters). Eluate from the SepPak cartridge was evaporated to dryness and resuspended in buffer A (20 mM ammonium hydroxide, pH 10) prior to fractionation by high pH reversed-phase chromatography using an Ultimate 3000 liquid chromatography system (Thermo). In brief, the sample was loaded onto an XBridge BEH C18 Column (130Å, 3.5 µm, 2.1 mm X 150 mm, Waters) in buffer A and peptides eluted with an increasing gradient of buffer B (20 mM Ammonium Hydroxide in acetonitrile, pH 10) from 0-95% over 60 min. The resulting fractions (6 in total) were evaporated to dryness and resuspended in 1% formic acid prior to analysis by nano-LC MSMS using an Orbitrap Fusion Lumos mass spectrometer (Thermo).

##### Nano-LC Mass Spectrometry

High pH RP fractions were further fractionated using an Ultimate 3000 nano-LC system in line with an Orbitrap Fusion Lumos mass spectrometer (Thermo). In brief, peptides in 1% (vol/vol) formic acid were injected onto an Acclaim PepMap C18 nano-trap column (Thermo). After washing with 0.5% (v/v) acetonitrile 0.1% (v/v) formic acid peptides were resolved on a 250 mm × 75 μm Acclaim PepMap C18 reverse phase analytical column (Thermo) over a 150 min organic gradient, using 7 gradient segments (1-6% solvent B over 1 min, 6-15% B over 58 min, 15-32% B over 58 min, 32-40% B over 5 min, 40-90% B over 1 min, held at 90% B for 6 min and then reduced to 1% B over 1 min) with a flow rate of 300 nl min^−1^. Solvent A was 0.1% formic acid and Solvent B was aqueous 80% acetonitrile in 0.1% formic acid. Peptides were ionized by nano-electrospray ionization at 2.0kV using a stainless-steel emitter with an internal diameter of 30 μm (Thermo) and a capillary temperature of 300°C.

All spectra were acquired using an Orbitrap Fusion Lumos mass spectrometer controlled by Xcalibur 3.0 software (Thermo) and operated in data-dependent acquisition mode using an SPS-MS3 workflow. FTMS1 spectra were collected at a resolution of 120 000, with an automatic gain control (AGC) target of 200 000 and a max injection time of 50 ms. Precursors were filtered with an intensity threshold of 5000, according to charge state (to include charge states 2-7) and with monoisotopic peak determination set to Peptide. Previously interrogated precursors were excluded using a dynamic window (60 s +/-10 ppm). The MS2 precursors were isolated with a quadrupole isolation window of 0.7 m/z. ITMS2 spectra were collected with an AGC target of 10 000, max injection time of 70 ms and CID collision energy of 35%.

For FTMS3 analysis, the Orbitrap was operated at 50 000 resolution with an AGC target of 50 000 and a max injection time of 105 ms. Precursors were fragmented by high energy collision dissociation (HCD) at a normalised collision energy of 60% to ensure maximal TMT reporter ion yield. Synchronous Precursor Selection (SPS) was enabled to include up to 10 MS2 fragment ions in the FTMS3 scan.

##### Data Analysis

The raw data files were processed and quantified using Proteome Discoverer software v2.4 (Thermo) and searched against the UniProt Human database (downloaded January 2021: 169297 entries) using the SEQUEST HT algorithm. Peptide precursor mass tolerance was set at 10 ppm, and MS/MS tolerance was set at 0.6 Da. Search criteria included oxidation of methionine (+15.995 Da), acetylation of the protein N-terminus (+42.011 Da) and Methionine loss plus acetylation of the protein N-terminus (−89.03 Da) as variable modifications and carbamidomethylation of cysteine (+57.0214) and the addition of the TMTpro mass tag (+304.207) to peptide N-termini and lysine as fixed modifications. Searches were performed with full tryptic digestion and a maximum of 2 missed cleavages were allowed. The reverse database search option was enabled and all data was filtered to satisfy false discovery rate (FDR) of 5%.

#### Light Microscopy Analysis

##### Fixed Cell Confocal Microscopy

For staining with dyes, cells on glass coverslips were incubated with 100 nM MitoTrackerRed CMXRos (Invitrogen) for 30 min at 37°C then washed twice in HBSS. Cells were then incubated with 5 µg/mL Wheat Germ Agglutin (WGA) Alexa Fluor 633 (Thermo) for 10 min at RT then washed three times in HBSS followed by fixation (protocol adapted from [62]). Coverslips were washed three times in HBSS then incubated in fixative 1 (2% (w/v) paraformaldehyde (PFA), 0.05% (w/v) glutaraldehyde, 0.2 M HEPES in PBS) for 20 min at RT. Fixative 1 was removed and coverslips were incubated with fixative 2 (4% (w/v) PFA, 0.2 M HEPES in PBS) for a further 20 min at RT. Coverslips were washed once with 100 mM glycine in PBS for 4 min followed by three quick washes in PBS. Coverslips were dipped in ddH_2_O before mounting using Fluoromount-G mounting medium (Thermo). Cells were imaged using a Leica confocal microscope (SP8) with ‘lightning’ adaptive image restoration, and Leica Application Suite X (LAS X) software platform. Laser lines used were 488, 562, and 633 nm, with gain set to allow maximum sensitivity without saturation, and z-stacks were taken.

For immunocytochemistry, following fixation and quenching (as above), cells were permeabilised by incubation in 0.1% (v/v) Triton-X in PBS for 5 min, washed in PBS and blocked with 3% (w/v) BSA in PBS for 30 min. Coverslips were incubated on a drop of primary antibody in 3% BSA overnight at 4°C. Coverslips were washed extensively in PBS and incubated with the appropriate secondary antibody (2 h, RT) in 3% BSA, then washed again. Coverslips were dipped in ddH2O and mounted using Fluoromount-G. Coverslips were left to dry overnight and imaged using a Leica confocal microscope (SP5II or SP8) and LAS X software platform. Laser lines used were 405, 488, 562, and 633 nm, with gain set to allow maximum sensitivity without saturation, and z-stacks were taken.

##### Live Cell Confocal Microscopy

Cells were seeded onto 35 mm glass bottomed dishes (Corning). Immediately prior to imaging, cells were washed in HBSS, transferred to imaging media (10% FBS, 2 mM L-Glutamine, 10 mM D-Galactose, 1 mM Sodium Pyruvate, 40 mM HEPES in phenol red free DMEM), and mitochondria stained with 25 nM TMRM for 30 min at 37°C prior to imaging (TMRM retained in buffer throughout imaging). Cells were imaged immediately using a Leica SP8 confocal microscope and LAS X software platform at 37°C, using 488 and 562 nm laser lines.

##### Live Cell Spinning Disk Microscopy

Cells were seeded onto 35 mm glass bottomed dishes (Corning). Immediately prior to imaging, cells were incubated with MitoTracker Green for 30 min at 37°C, washed three times in HBSS and transferred into imaging media (as above). Imaging was performed using an Olympus IXplore SpinSR system at 37°C, using 488 and 561 nm laser lines. A z-stack was taken with 10 z-slices, taken at regular intervals (see figure legend).

##### Live Cell Structured Illumination Microscopy

Transfected cells were grown to confluency on glass coverslips prior to staining with 100 nM MitoTracker Green^TM^ for 30 min at 37°C in complete growth medium. Coverslips were washed in HBSS and incubated in complete growth medium in Attofluor™ Cell Chambers (Invitrogen) for live imaging using super-resolution 3D structured illumination microscopy (SR-3D SIM) on an OMX v4 microscope (GE Healthcare) at 37°C, using 488 and 568 nm laser lines and a 60x objective.

##### Light Microscopy Image Analysis

All image analysis was performed using the FiJi image processing package [63, 64]. Macros were written by Dr Stephen Cross (Wolfson Bioimaging Facility, University of Bristol), within the FiJi plugin Modular Image Analysis (MIA) package version 0.21.0, which is publicly accessible on GitHub with a linked version specific DOI from Zenodo [65]. Where data was analysed manually, data was randomised/blinded prior to manual analysis.

Mitochondrial pre-processing, as well as branch and network analysis was carried out as described previously [66, 67], using a FiJi plugin adapted from the mitochondrial network analysis (MiNA) toolset [68]. Briefly, for pre-processing, cells of interest were outlined and outside cleared, prior to z-stack max intensity projection, contrast enhancement and background subtraction. Median, unsharp, and tubeness filters were applied. For branch/network analysis, images were binarised and skeletonised, and the skeleton analysed to obtain information on the branches and pixels, allowing analysis of mitochondrial branching as a readout of mitochondrial network complexity and fragmentation.

Analysis of mitochondrial width was carried out by manual measurement. Data was blinded and the same region of mitochondria was measured in each cell (100 x 100 pixels at 2 ‘o’ clock relative to the nucleus) to maintain unbiased results. Five measurements were taken across each mitochondrion (parallel to cristae) and averaged to obtain an average width for the given mitochondrion. An equal number of mitochondria were analysed for each condition.

Circularity analysis was carried out using a macro that classifies mitochondria based on their shape, providing a circularity index, where a value of 1 represents a perfect circle.

Aggregation analysis was carried out using a macro, on images acquired by SIM. Mitochondria (channel 1) are detected in 2D within a region of interest (ROI, *i.e.,* a transfected cell) and the pixel intensity of the trapping substrate (channel 2) is measured as a 4-pixel wide strip at the edge of the mitochondrion. The background channel 2 intensity (intensity at 10-14 pixels from the edge of the mitochondrion) is subtracted from the raw intensity surrounding the mitochondrion, to account for differing expression levels between cells. This provides values for the intensity directly surrounding mitochondria, and after thresholding, these values highlight the proportion of mitochondria within a cell or population of cells that have trapped protein on their outside.

#### NanoLuc Assay

NanoLuc assays were carried out exactly as described previously [21]. Briefly, cells on standard white flat-bottom 96-well plates were washed with HBSS and incubated in HBSS imaging buffer (HBSS supplemented with 5 mM D-(+)-glucose,10 mM HEPES (Santa Cruz Biotechnology, Germany), 1 mM MgCl_2_, 1 mM CaCl_2_; pH 7.4). A fluorescence read was taken at 605/670 nm using monochromators with gain set to allow maximum sensitivity without saturation, using a BioTek Synergy Neo2 plate reader (Agilent, UK). Cells were transferred to permeabilised cell assay master mix (225 mM mannitol, 10 mM HEPES, 2.5 mM MgCl_2_, 40 mM KCl, 2.5 mM KH_2_PO_4_, 0.5 mM EGTA; pH 7.4) supplemented with 5 mM succinate, 1 µM rotenone, 0.1 mg/mL creatine kinase, 5 mM creatine phosphate, 1 mM ATP, 0.1% (v/v) Prionex, 3 nM rPFO (purified in house), 20 µM GST-Dark (purified in house), 1:800 furimazine (Nano-Glo® Luciferase Assay System; Promega). A baseline read of 30 seconds of background luminescent signal was taken prior to injection of purified substrate protein (*Su9-EGFP-pep86*) to 1 µM final concentration, followed by a further bioluminescence read corresponding to import, lasting 30 min. Bioluminescence was read using a BioTek Synergy Neo2 plate reader or a CLARIOstar Plus plate reader (BMG LabTech, UK) without emission filters with gain set to allow maximum sensitivity without saturation, and with acquisition time of 0.1 sec per well. Row mode was used, and reads were taken every 6 sec or less, with wells in triplicate.

#### Statistical Analysis

Statistical significance between groups was determined using unpaired Student’s t-Test or one and two-way ANOVA with interaction if more than two groups were analysed. ANOVA was nested if multiple objects were analysed per biological replicate. Following ANOVA, p values were adjusted for multiple comparisons through Tukey’s *post-hoc* test and differences were considered significant at 5% level. Statistical analyses were performed using Graph Pad Prism version 9 (GraphPad Software, Inc., San Diego, CA, USA).

**Fig. S1.**
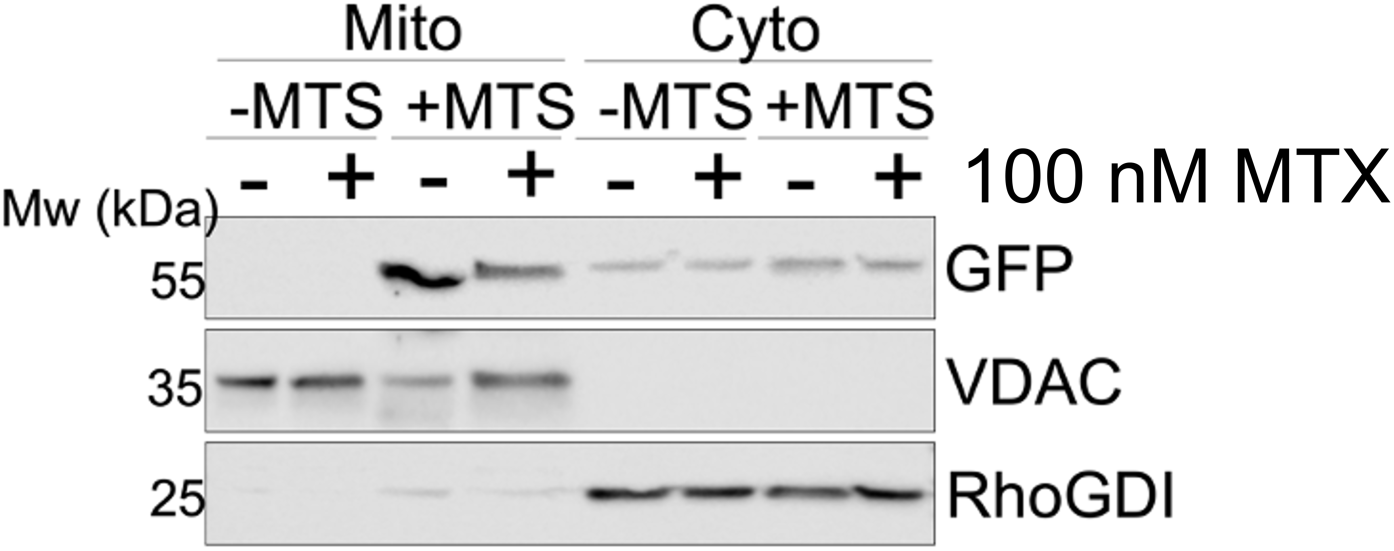
Representative Western blot showing expression (48 h) of *EGFP-DHFR* (-MTS) or *Su9-EGFP-DHFR* (+MTS) in the cytosolic and crude mitochondrial fractions of HeLaGAL cells in the presence or absence of 100 nM MTX (48 h). The abundance of *EGFP-DHFR* or *Su9-EGFP-DHFR* in each fraction was visualised by probing with an anti-GFP antibody (Sigma; G1544). VDAC (Invitrogen; PA1-954A) was used as a mitochondrial loading control and RhoGDI (Abcam; ab133248) was used as a cytosolic loading control. N=4 biological replicates.

**Fig. S2.**
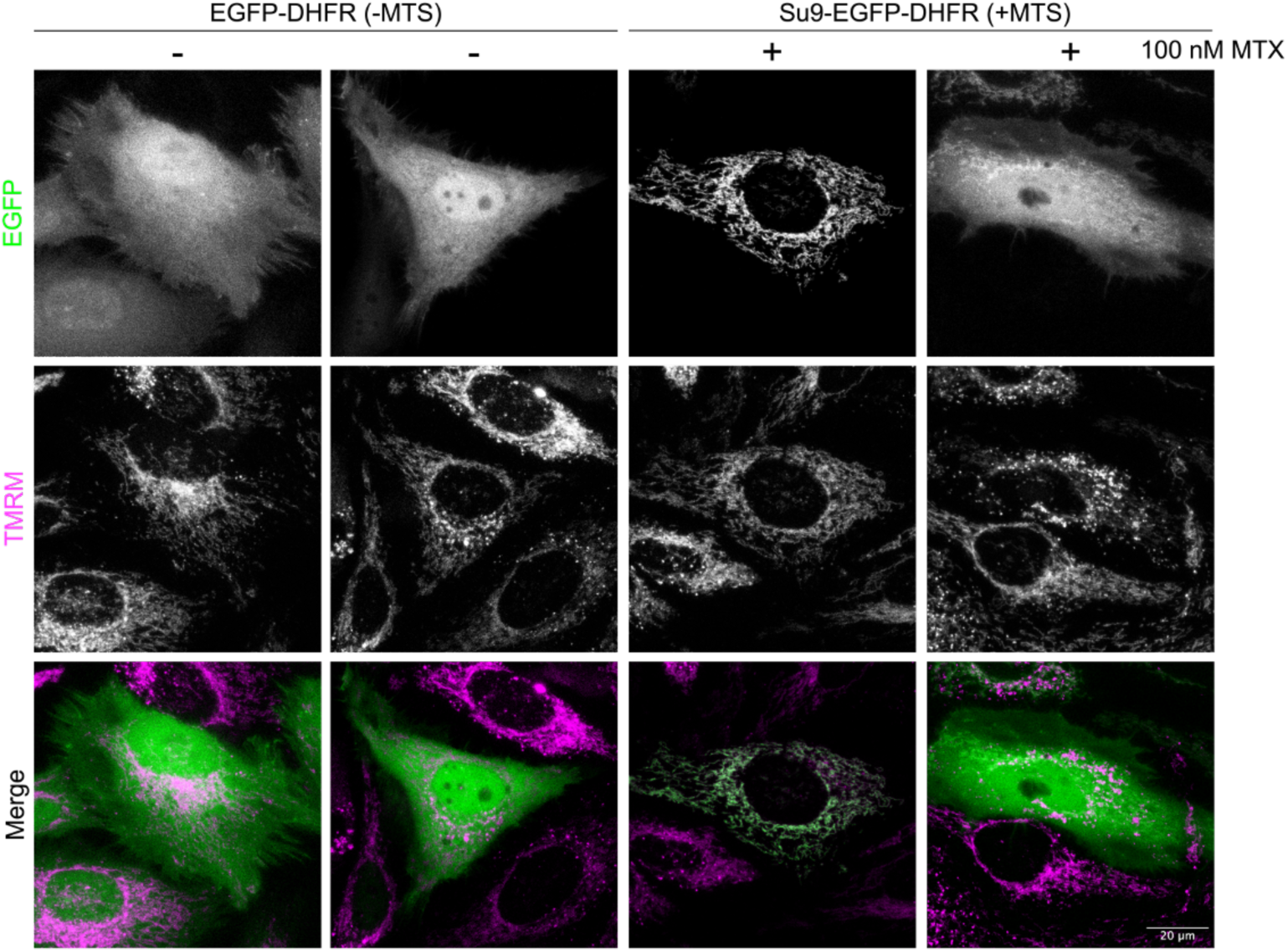
Representative confocal images of HeLaGAL cells treated with 100 nM MTX (or with vehicle; DMSO only) overnight prior to expression of *EGFP-DHFR* (-MTS; green) or *Su9-EGFP-DHFR* (+MTS; green) for 48 hours. Mitochondria were stained with 25 nM TMRM (magenta). Overlay shows cellular *EGFP-DHFR* or *Su9-EGFP-DHFR* localisation. N=3 biological replicates.

**Fig. S3.**
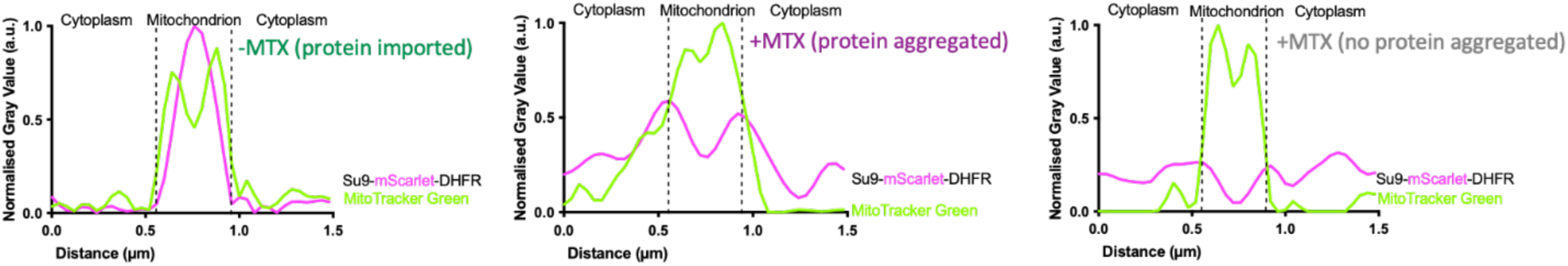
Representative plots of average pixel intensity across a mitochondrion in cells subjected to the trapping insult with (middle panel) or without (right) associated trapped precursor, or in cells not treated with MTX (with imported precursor; left panel).

**Fig. S4.**
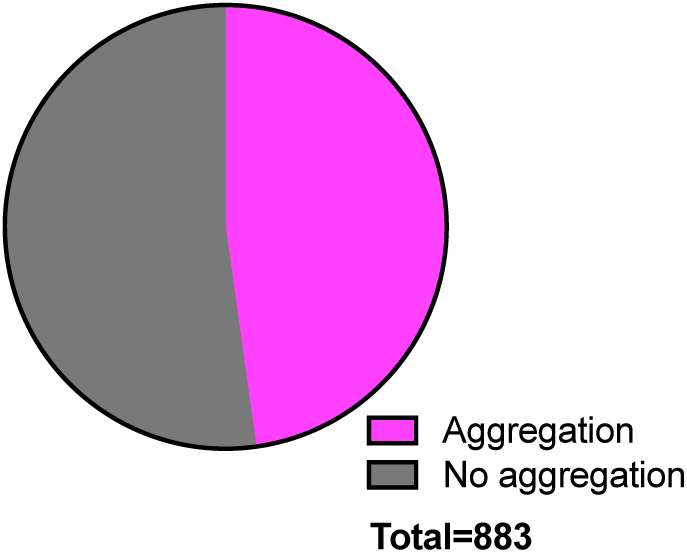
Quantification of the proportion of mitochondria per cell with (magenta) or without (grey) trapped precursor surrounding them. Analysis was done using a FiJi plugin as described in methods. N=3 biological replicates. Analysis included a total of 883 mitochondria.

**Fig. S5.**
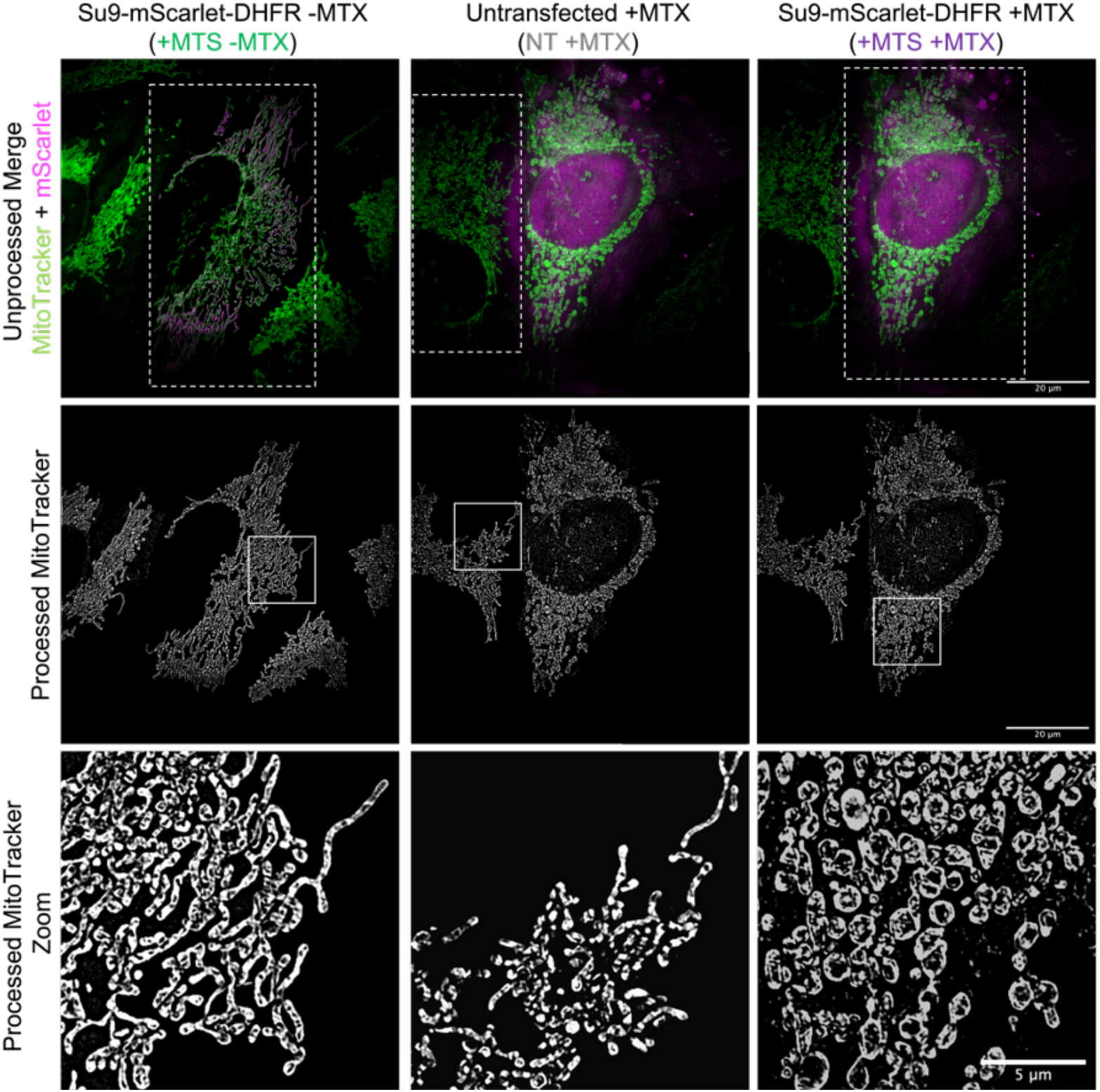
Representative SR 3D-SIM images showing mitochondria in HeLaGAL cells subjected to trapping (+MTS +MTX) or not (+MTS-MTX). Untransfected cells in the same well as cells subjected to trapping were used as an internal control for MTX treatment (NT +MTX). The top row shows a merge of MitoTracker Green (green) and mScarlet (magenta) prior to pre-processing. The middle row shows mitochondria (MitoTracker Green channel only) after pre-processing (described in methods). The bottom panel shows a zoom of an area of mitochondria (highlighted in the middle panel) to give a clearer view of mitochondrial morphology. N=3 biological repeats.

**Fig. S6.**
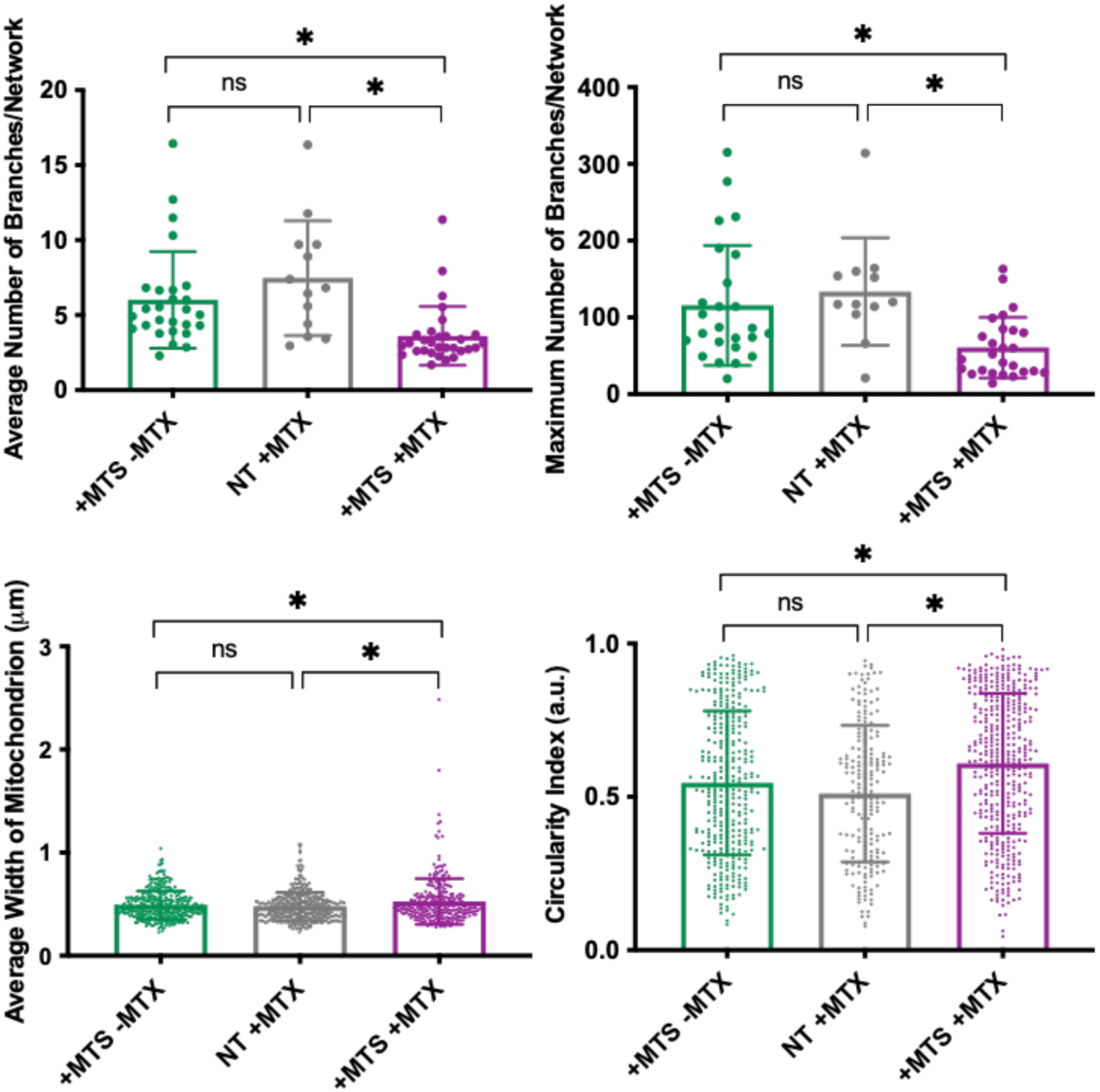
Quantification of mitochondrial morphology in terms of branching, width, and circularity. This was carried out for cells expressing *Su9-mScarlet-DHFR* in the absence of MTX (imported precursor; +MTS-MTX; green), internal control cells (untransfected but in the presence of 100 nM MTX; NT +MTX; grey), and transfected cells in the presence of MTX (trapped precursor; +MTS +MTX; purple). For *branching*, each point represents an individual cell from a separate field of view. N=27, 13, 30 cells for different conditions, respectively, taken from 3 independent biological repeats (N=3). For *width*, each point represents an individual mitochondrion, as an average of 5 measurements taken for any one mitochondrion, from a region from a selected field of view (independent cell). N=360, 470, 315 mitochondria, from 20 cells for each condition, respectively, taken from 3 independent biological repeats (N=3). For *circularity*, each point represents an individual mitochondrion. N=359, 199, 434 mitochondria, respectively, taken from 3 independent biological repeats (N=3). Error bars show SD. Nested one-way ANOVA and Tukey’s *post-hoc* test were used to determine significance.

**Fig. S7.**
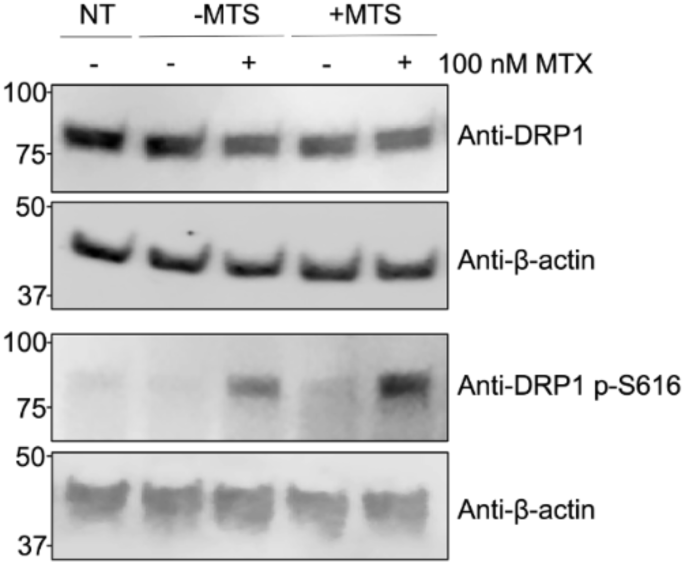
Representative Western blots showing abundance of total DRP1 (BD Biosciences; 6111113) or phospho-DRP1 S616 (CST; 4494) in whole cell lysates from HeLa cells expressing *EGFP-DHFR* or *Su9-EGFP-DHFR*, or untransduced (NT) cells in the absence of presence of 100 nM MTX. β-actin (Sigma; A2228) was used as a loading control. N=3 biological replicates.

**Fig. S8.**
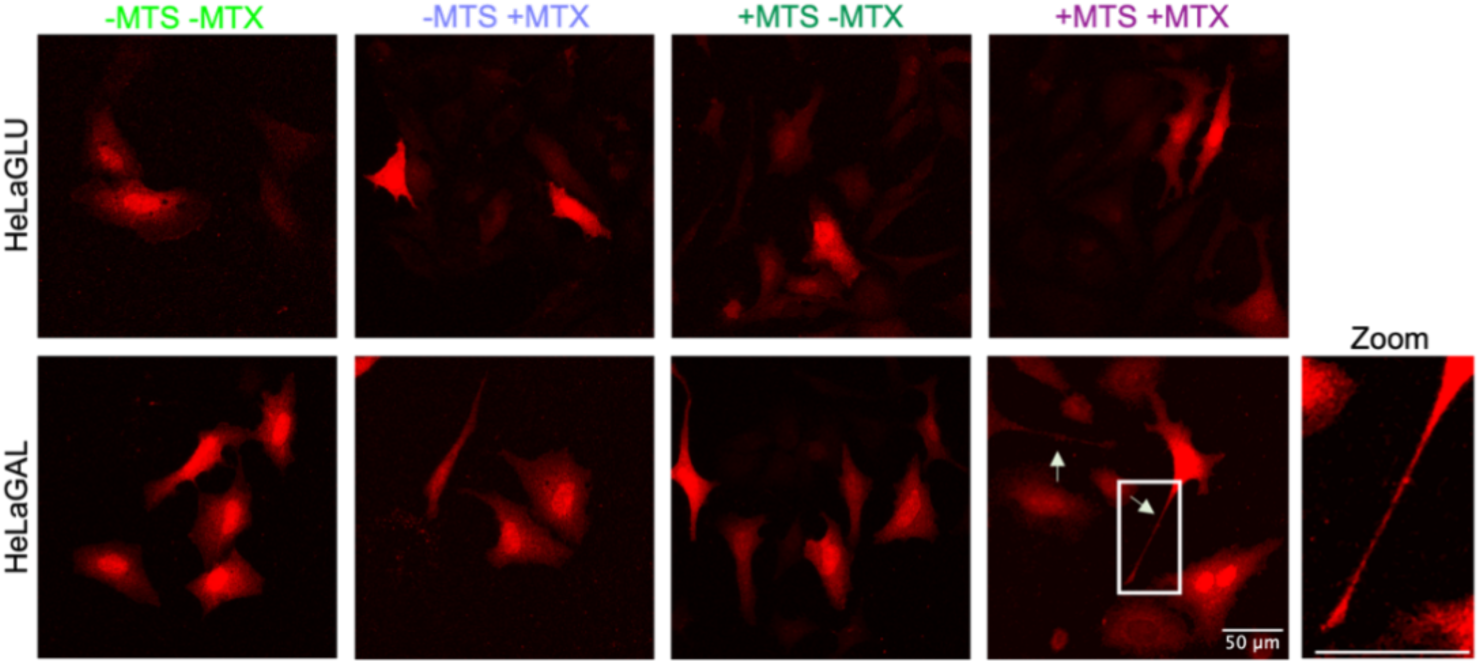
Representative confocal images showing morphology of HeLaGLU or HeLaGAL cells subjected to expression of *mCherry* (red) as well as *EGFP-DHFR* (-MTS) or *Su9-EGFP-DHFR* (+MTS) in the absence or presence of 100 nM MTX. Arrows point to protrusions, and box highlights region of interest displayed in zoom. N=4 biological replicates.

**Fig. S9.**
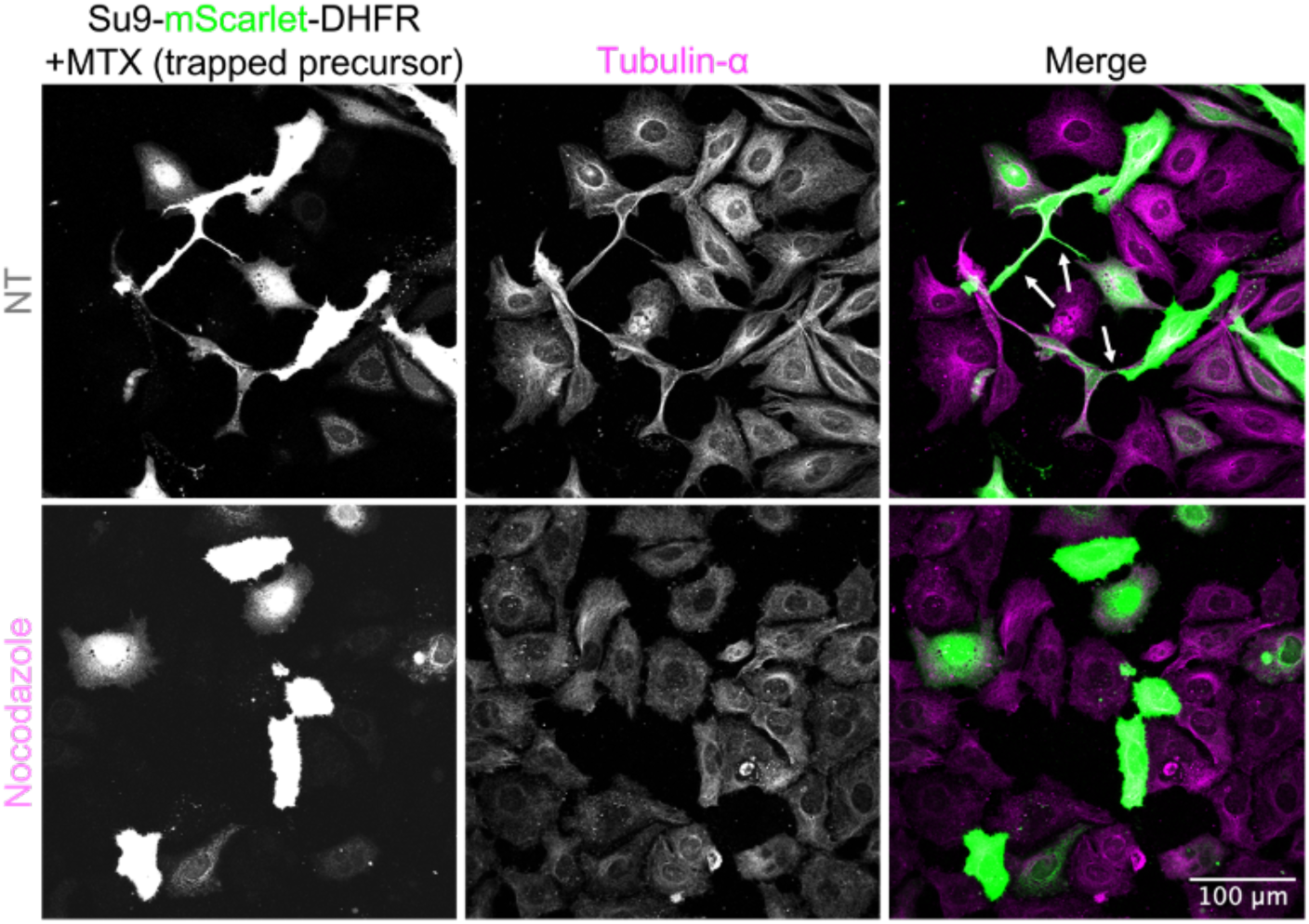
Representative confocal images of HeLaGAL cells exposed to precursor trapping in the absence or presence of TNT inhibitor nocodazole. HeLaGAL cells were subjected to trapping (*Su9-mScarlet-DHFR* +100 nM MTX; green) and either untreated (NT; DMSO only) or treated with 100 nM Nocodazole for 48 h (duration of precursor trapping). Cells were fixed and stained for tubulin-α (magenta; BioRad MCA78G) to visualise TNTs. N=3 biological replicates.

**Fig. S10.**
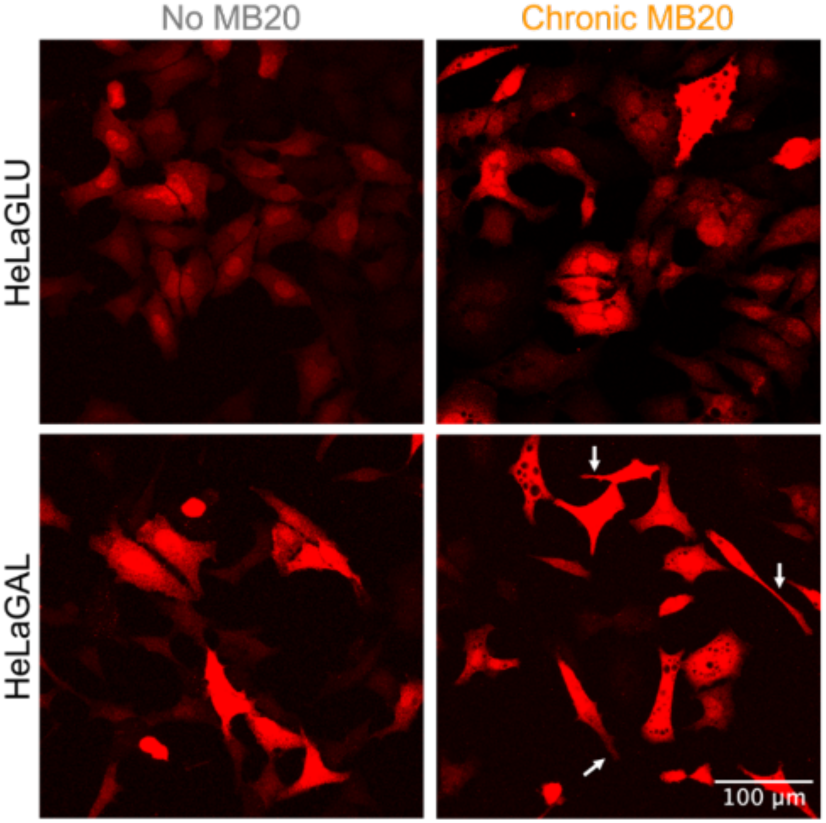
Representative images demonstrating morphology (shown by overexpressed *mCherry* cytosolic marker; red) of HeLaGLU or HeLaGAL cells treated with 10 µM MB20 for 48 h, visualised by confocal microscopy. Arrows highlight TNTs. N=6 biological replicates.

**Fig. S11.**
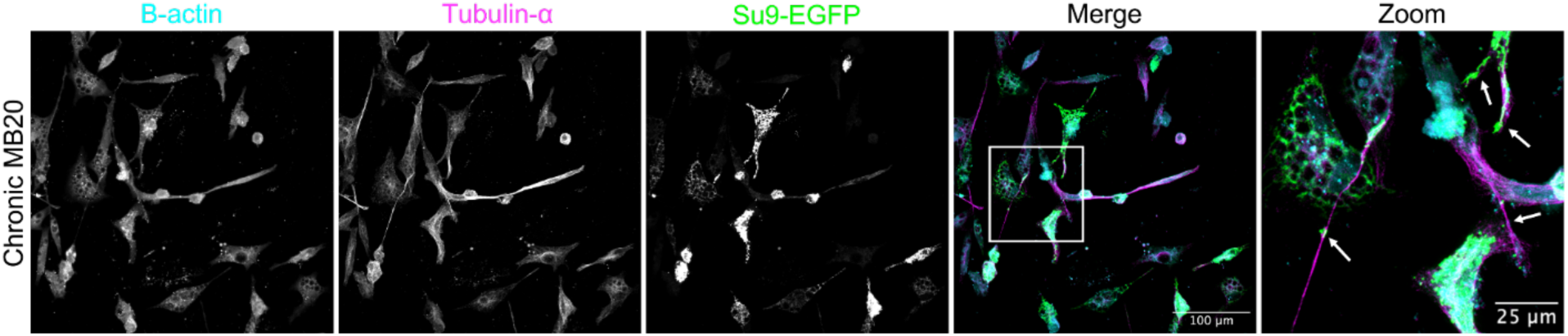
Representative images showing composition of MB20-induced protrusions. Cells expressing *Su9-EGFP* (mitochondrial marker; green) were treated for 48 h with 10 µM MB20. Fixed cells were stained for β-actin (cyan; Sigma A2228) and tubulin-α (magenta; BioRad MCA78G). Box shows zoom area and arrows in zoom highlight TNTs. N=3 biological replicates.

**Fig. S12.**
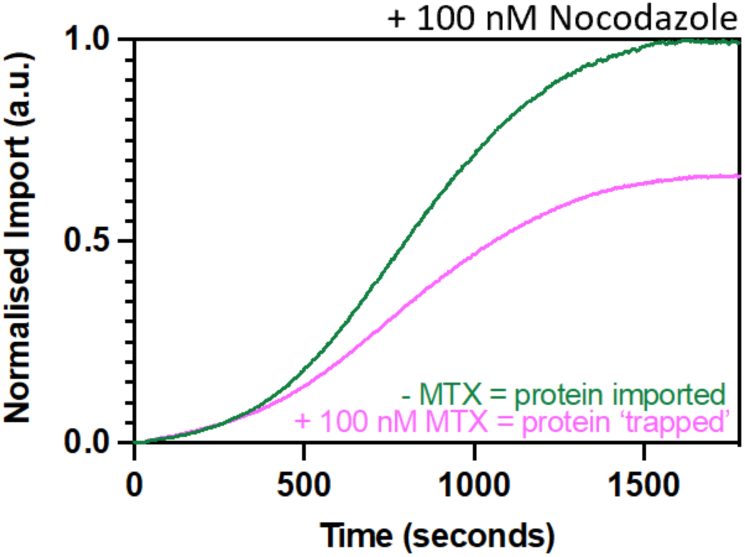
NanoLuc import trace showing the impact of TNT inhibitor nocodazole on import of a precursor protein in HeLaGAL cells subjected to precursor trapping. Cells were subjected to trapping (+MTX; pink) or not (-MTX; green) in the presence of 100 nM Nocodazole for 48 h prior to monitoring the import of *Su9-EGFP-pep86* by NanoLuc import assays. N=3 biological replicates, each with n=3 technical replicates.

**Fig. S13.**
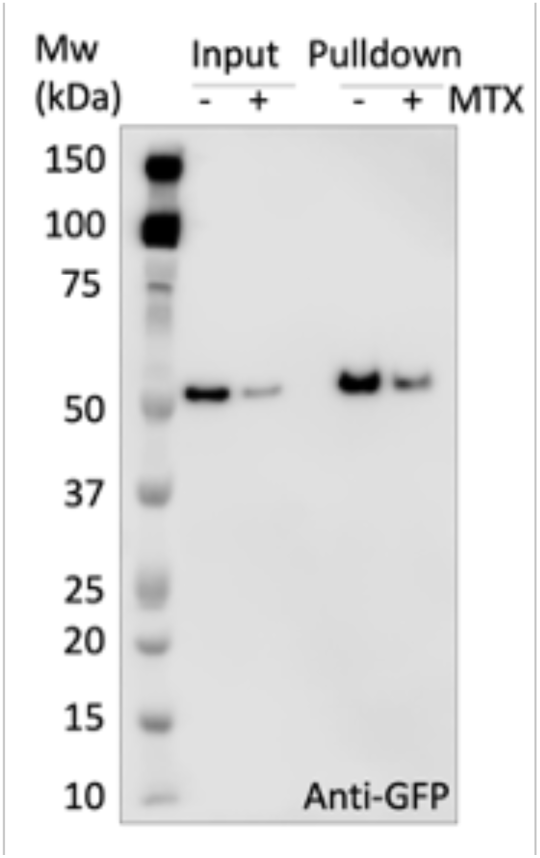
Representative Western blot showing GFP-tagged protein in the input and pulldown samples. Samples were prepared as described in schematic in Fig. 4D and probed against a GFP antibody (Sigma; G1544) to validate IP. N=3 biological replicates.

**Fig. S14.**
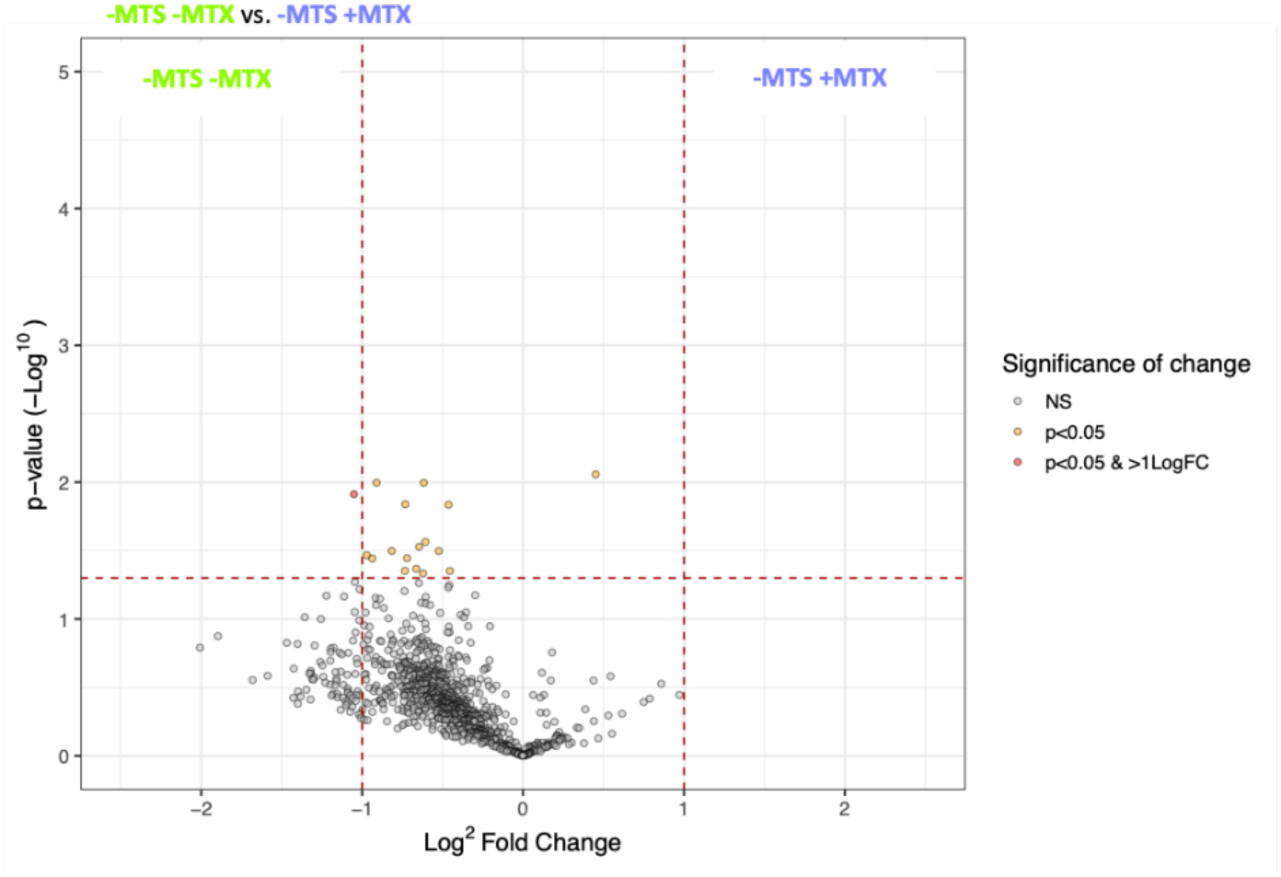
Volcano plot highlighting proteins enhanced in pulldown samples from mitochondria of cells overexpressing *EGFP-DHFR* (-MTS) in the presence or absence of 100 nM MTX, to control for background effect of MTX. N=3 biological replicates.

**Table S1.**
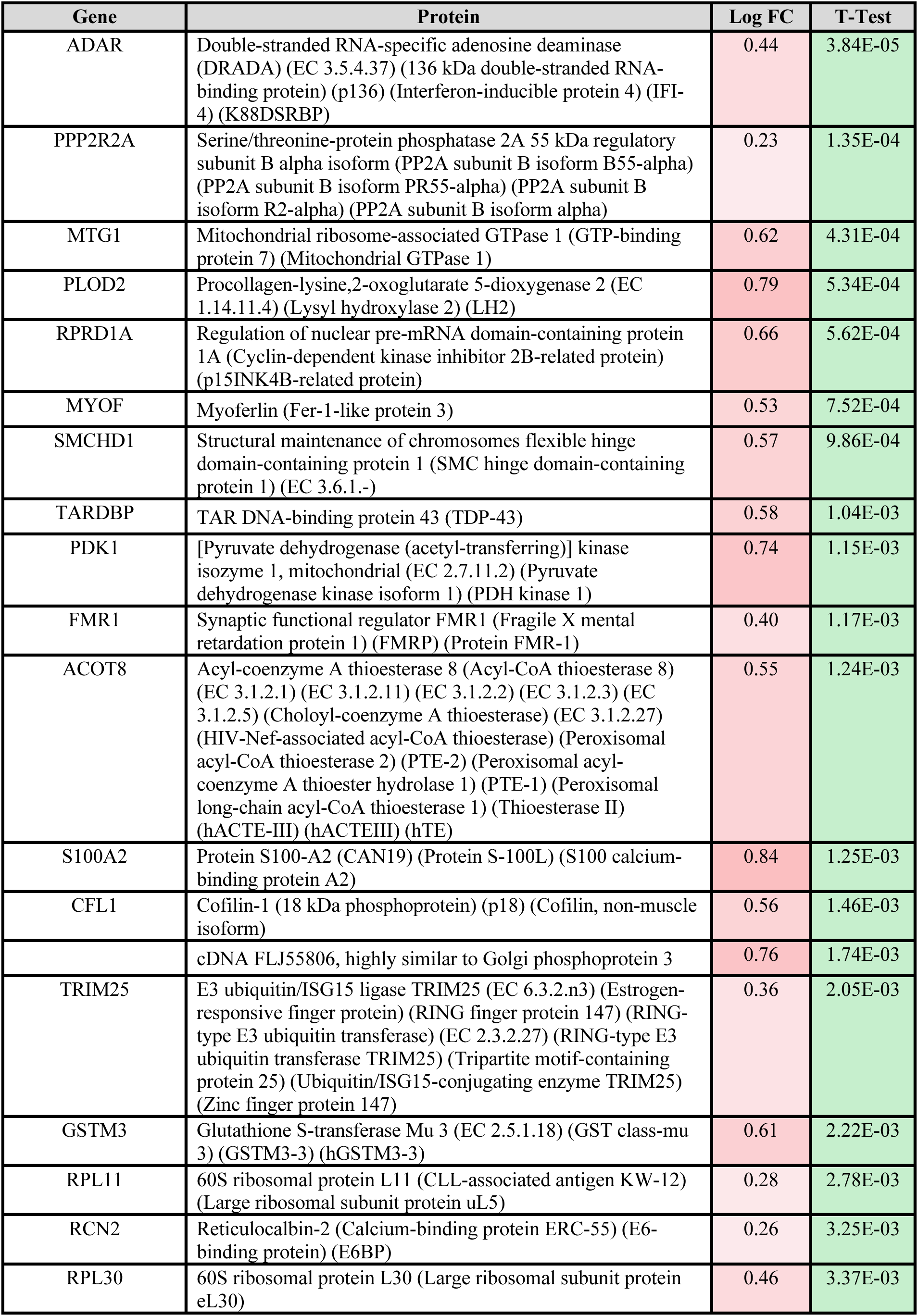
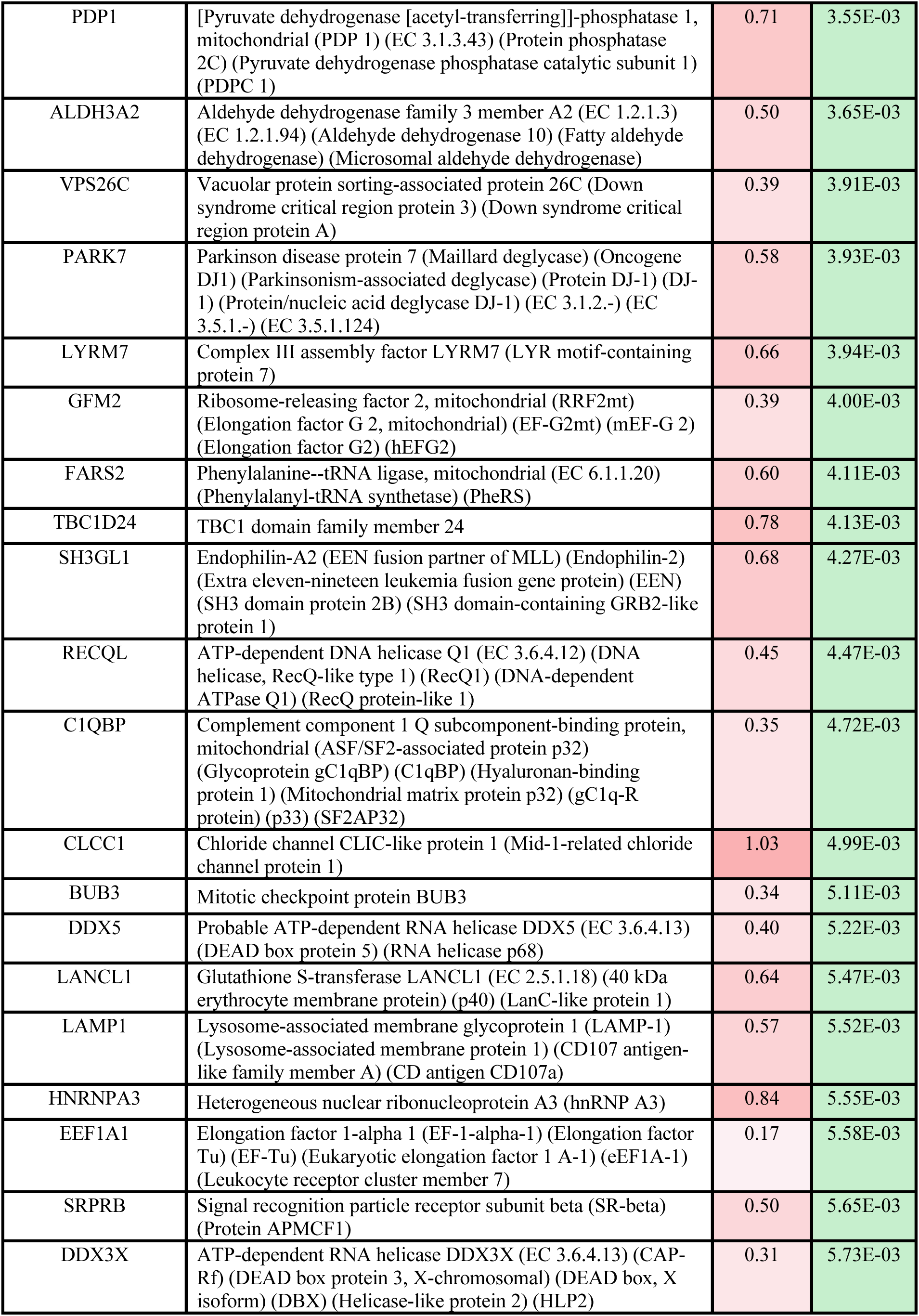
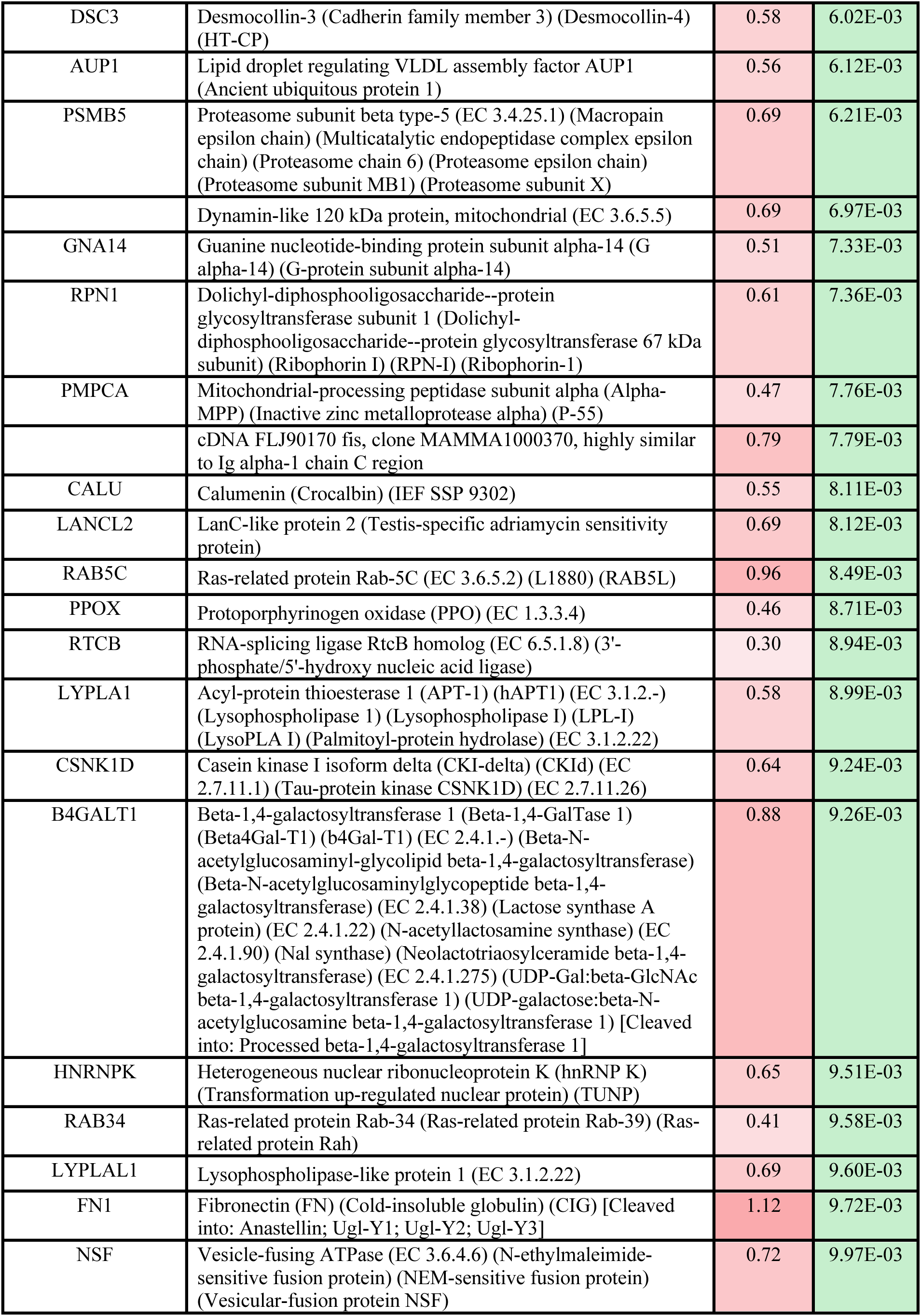
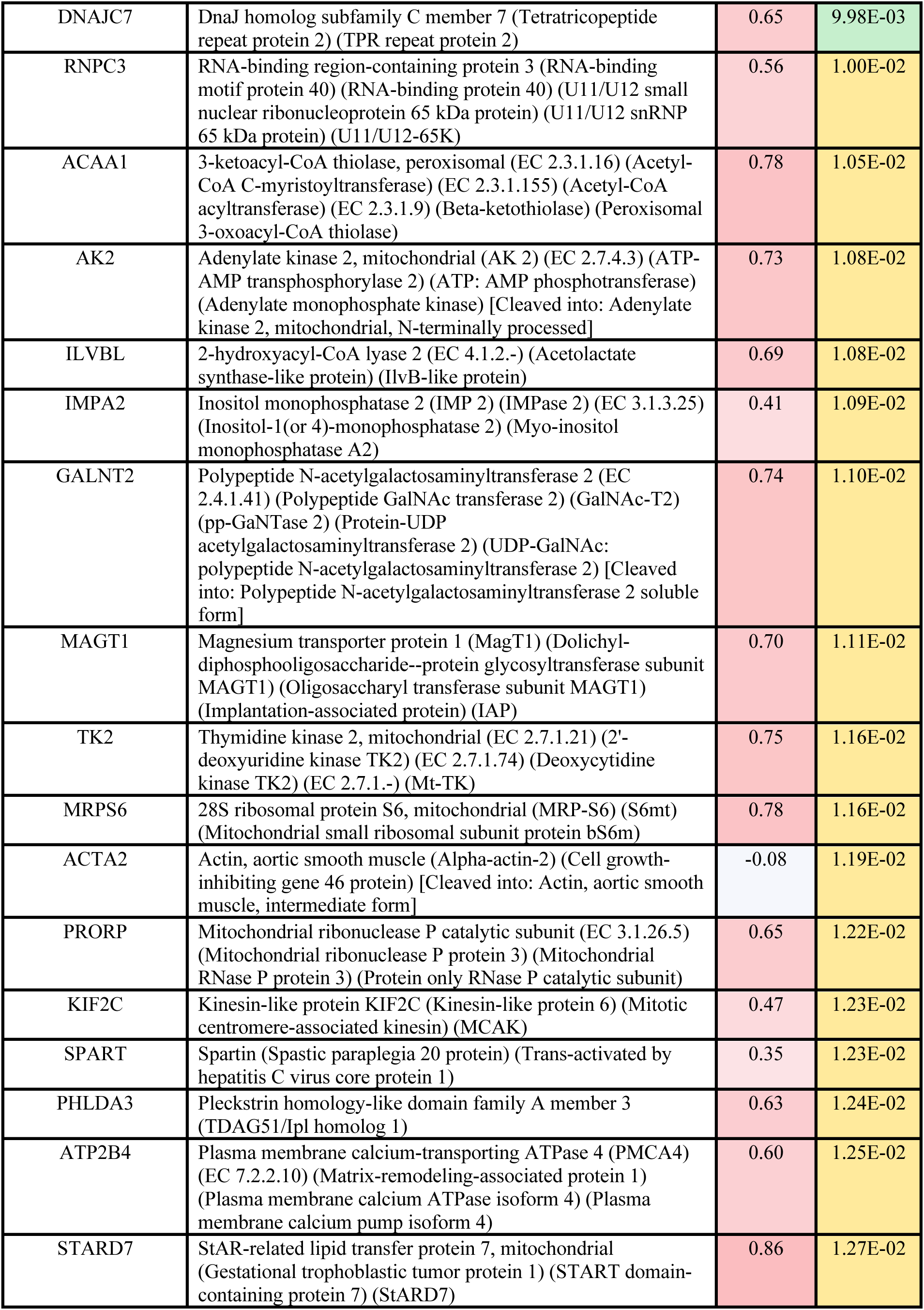
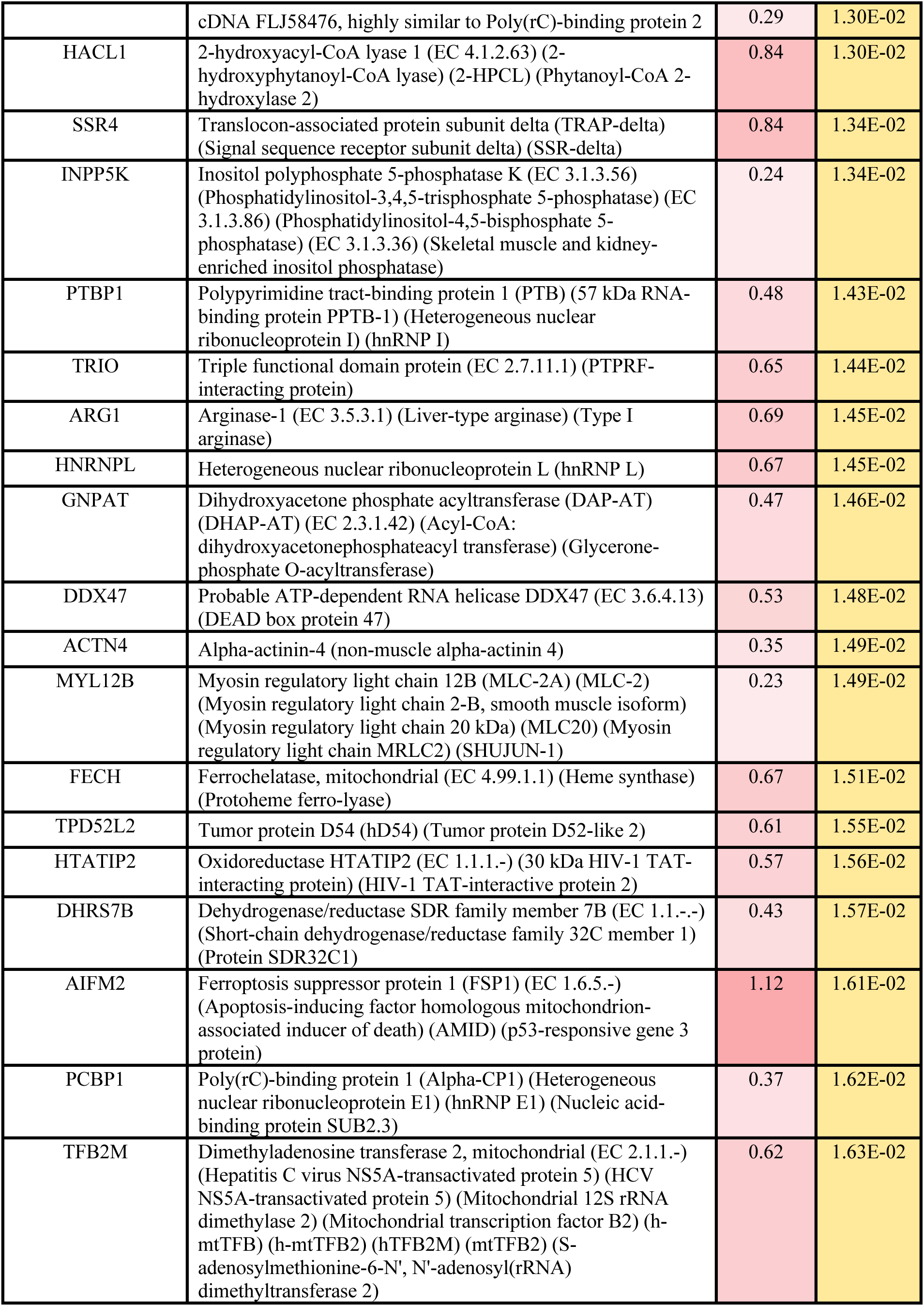
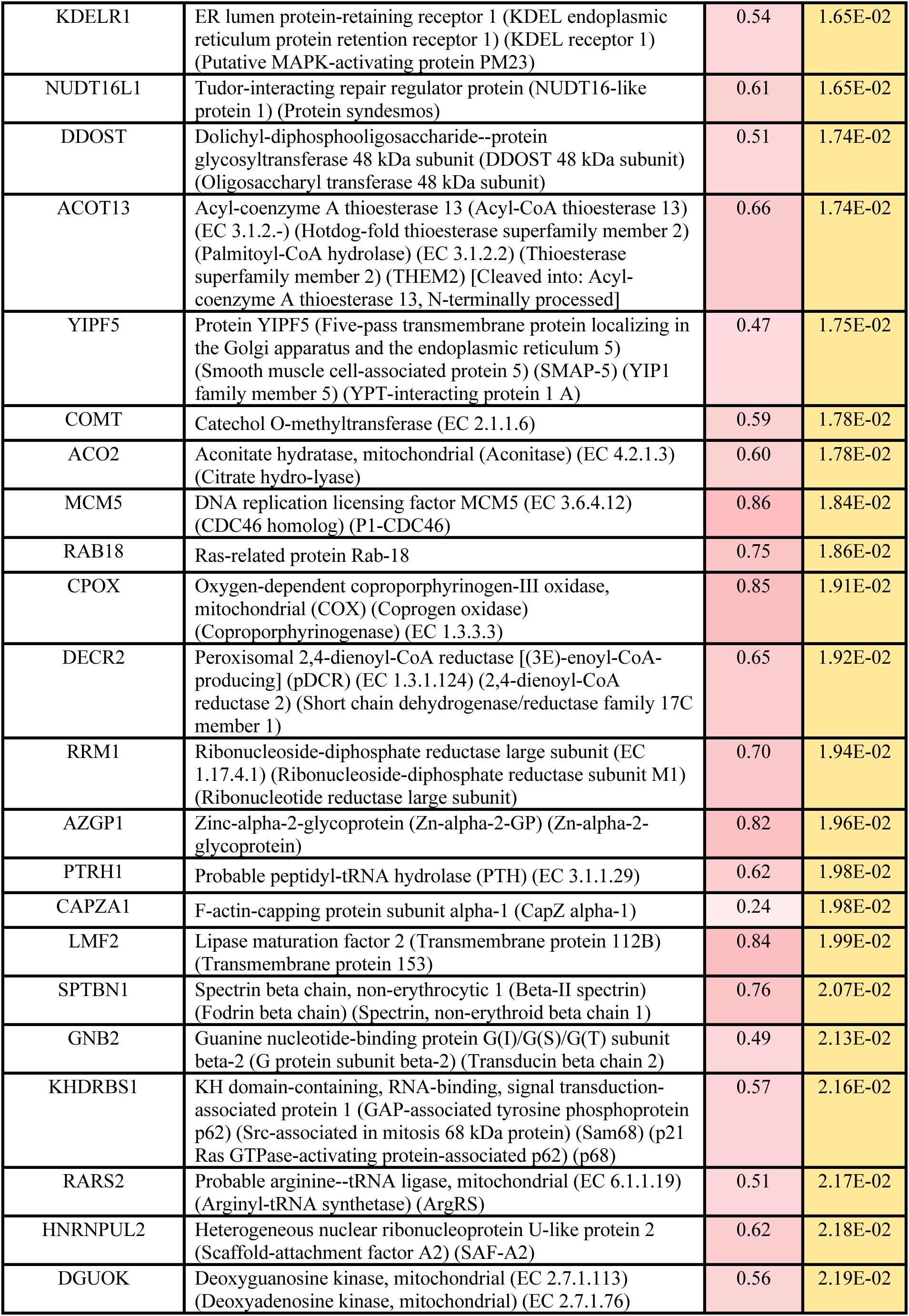
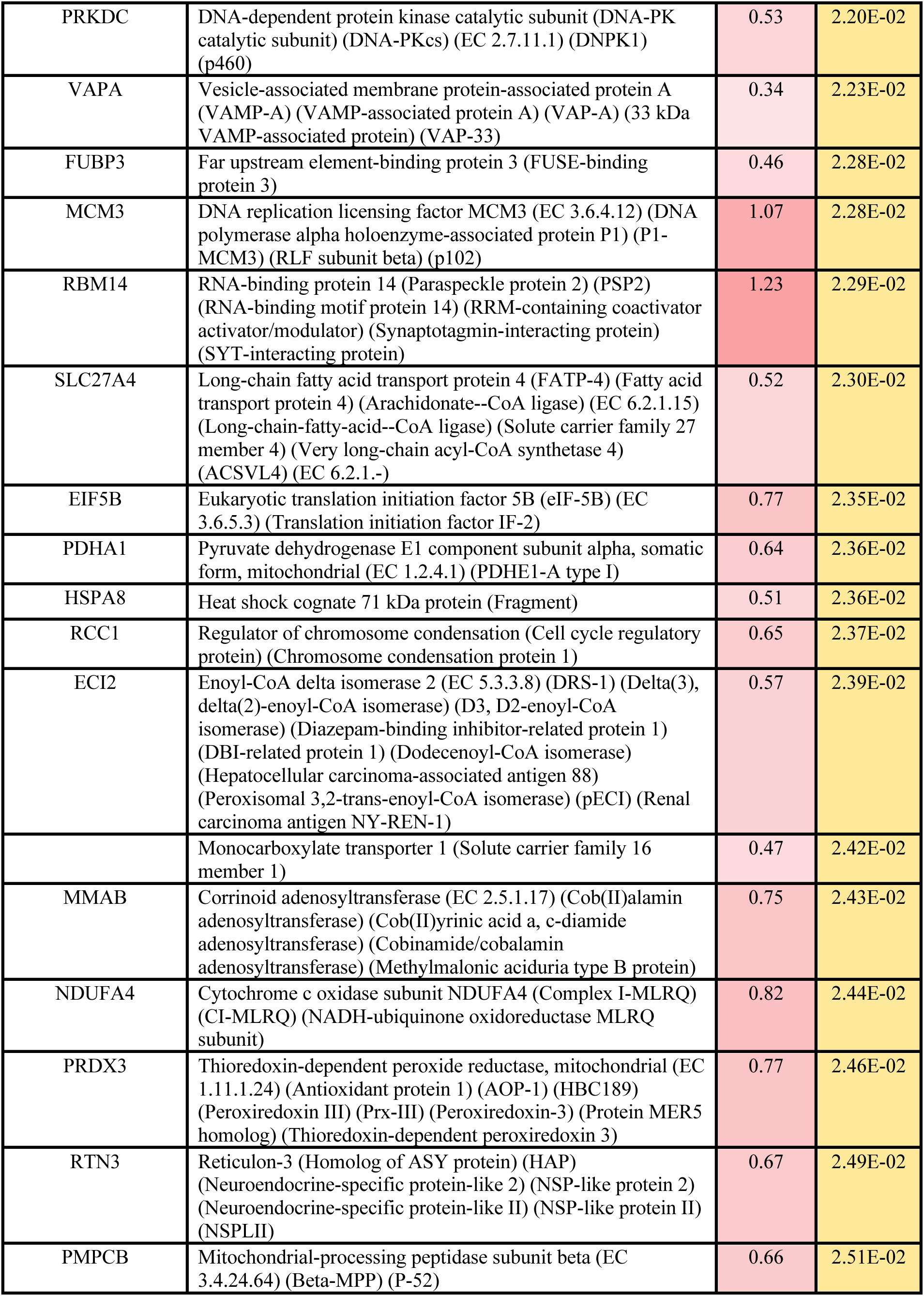
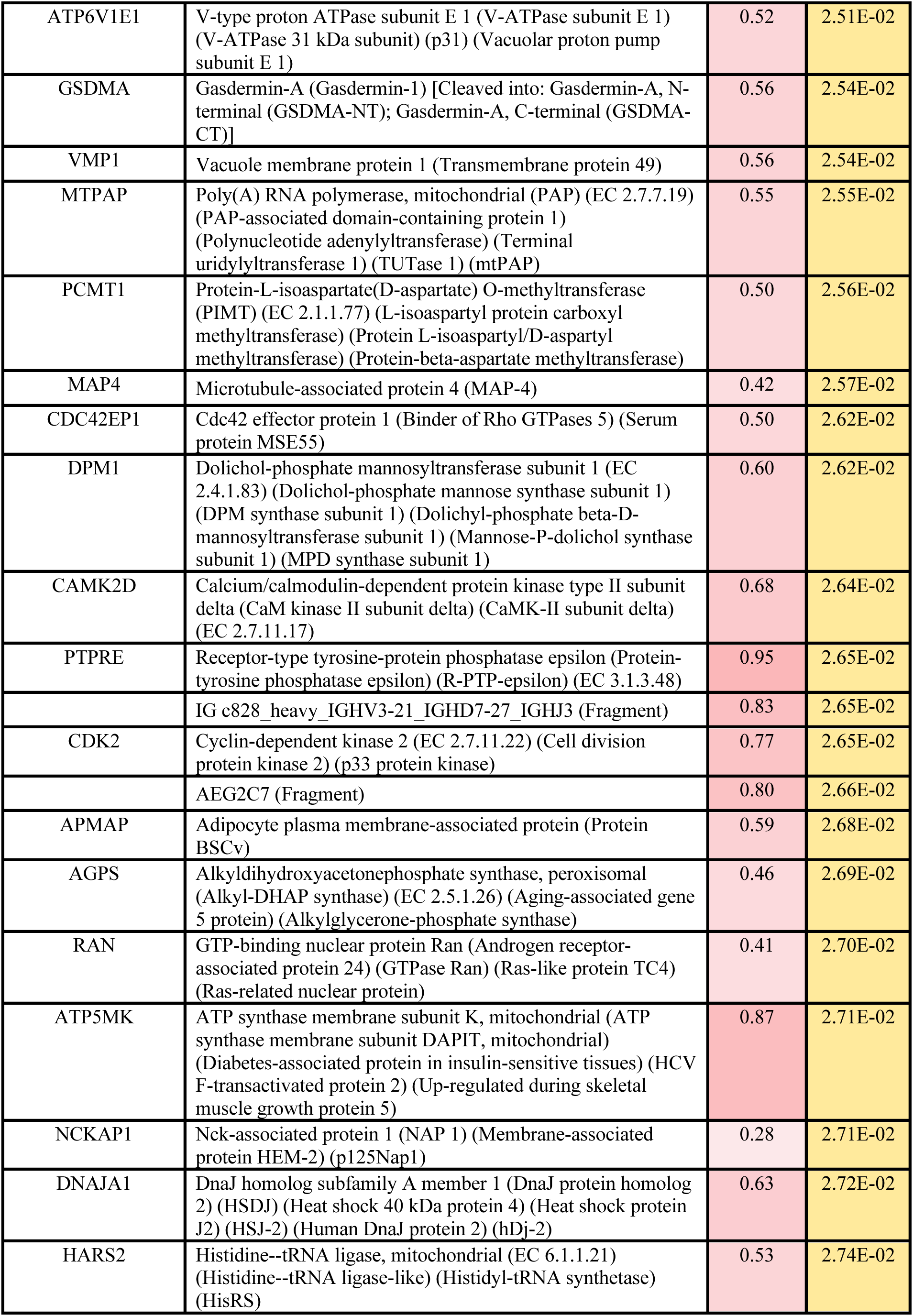
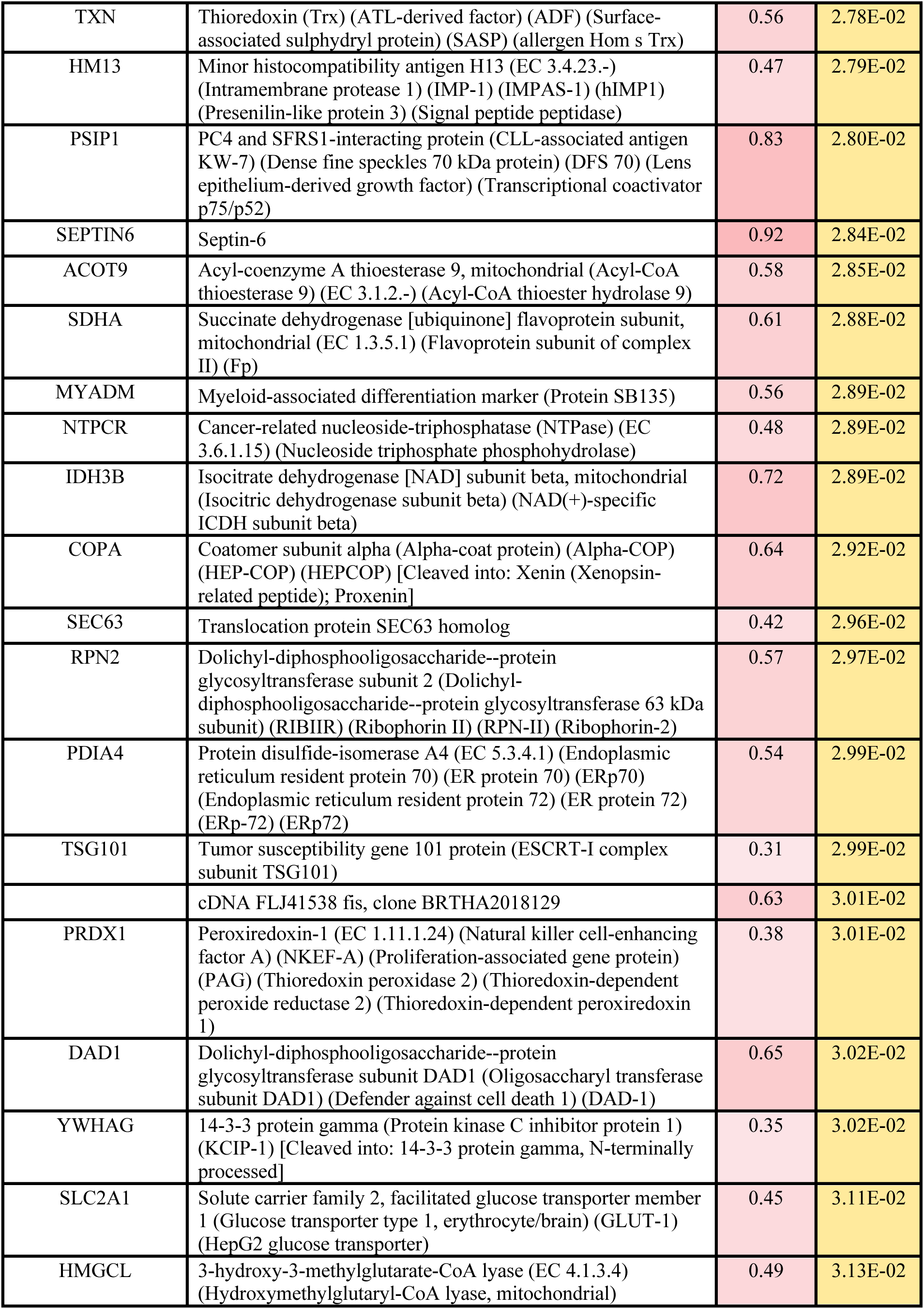
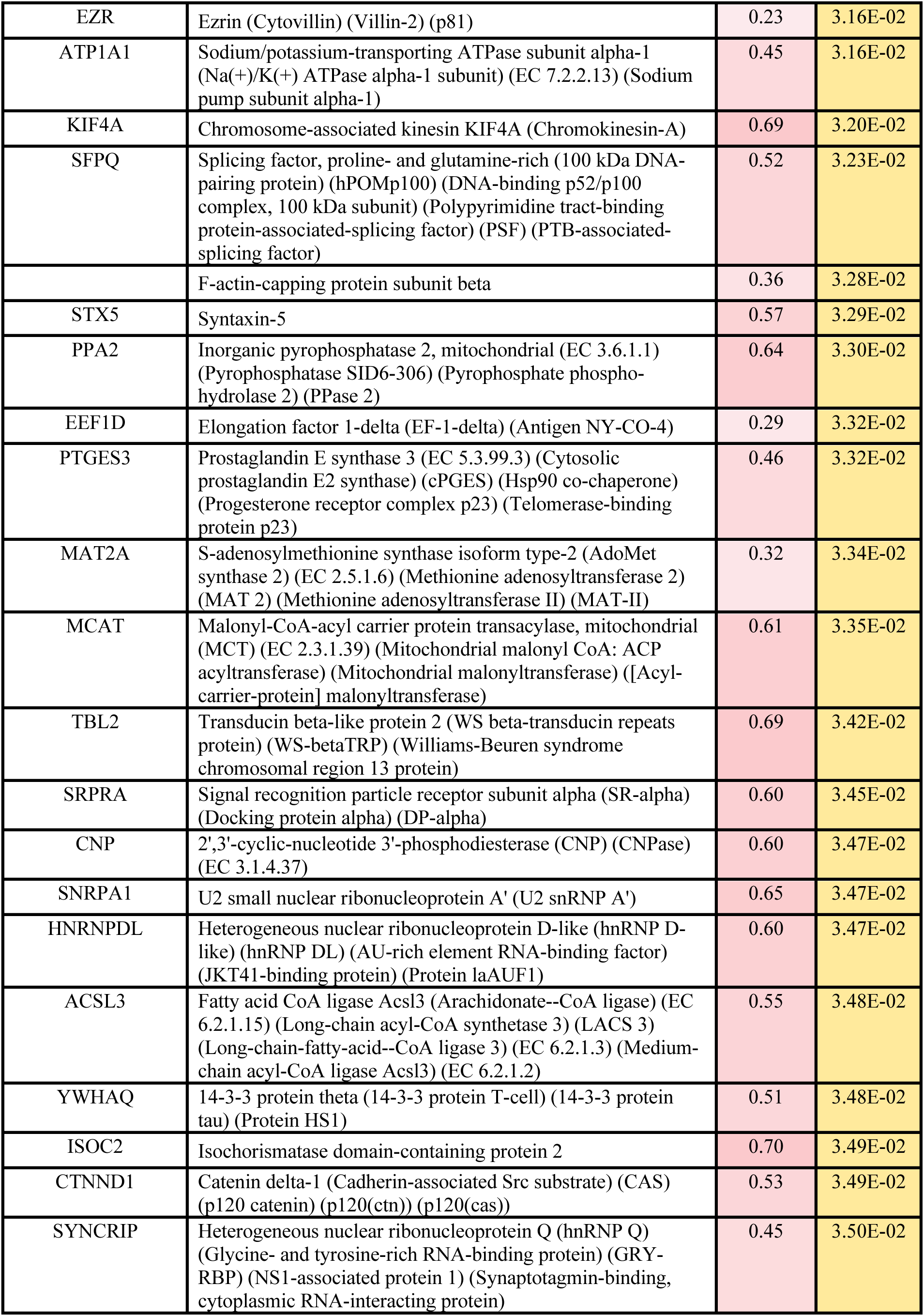
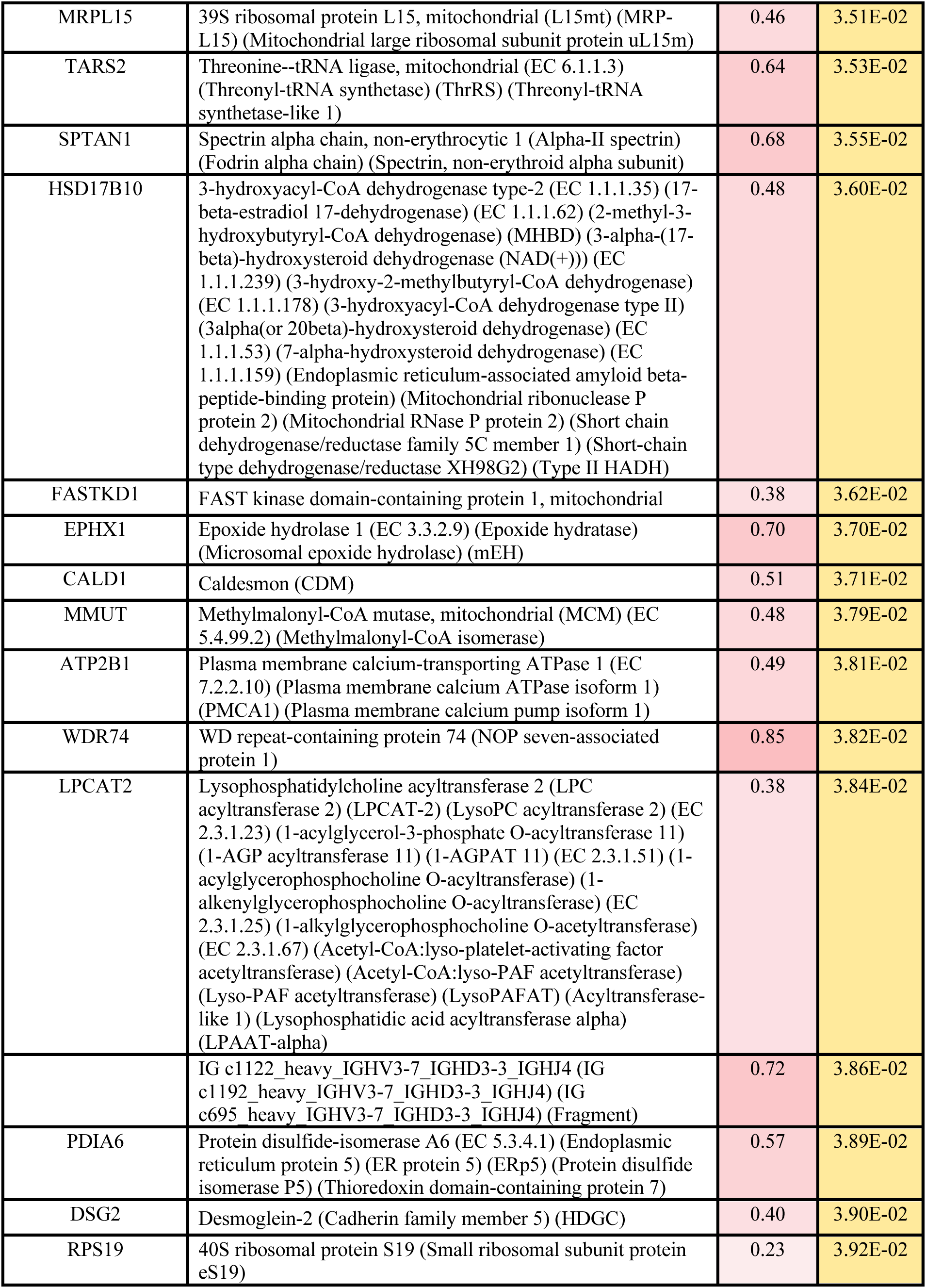
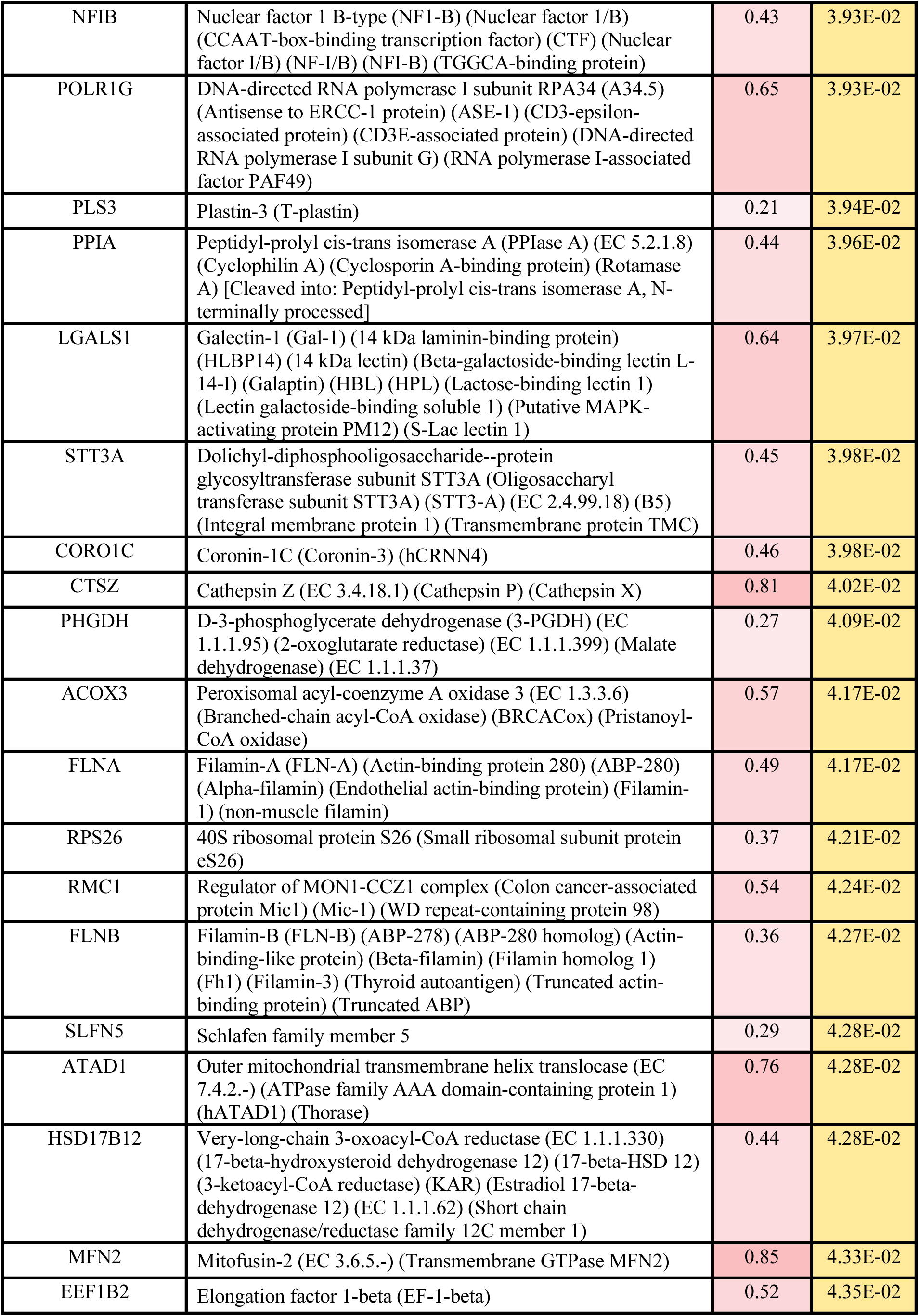
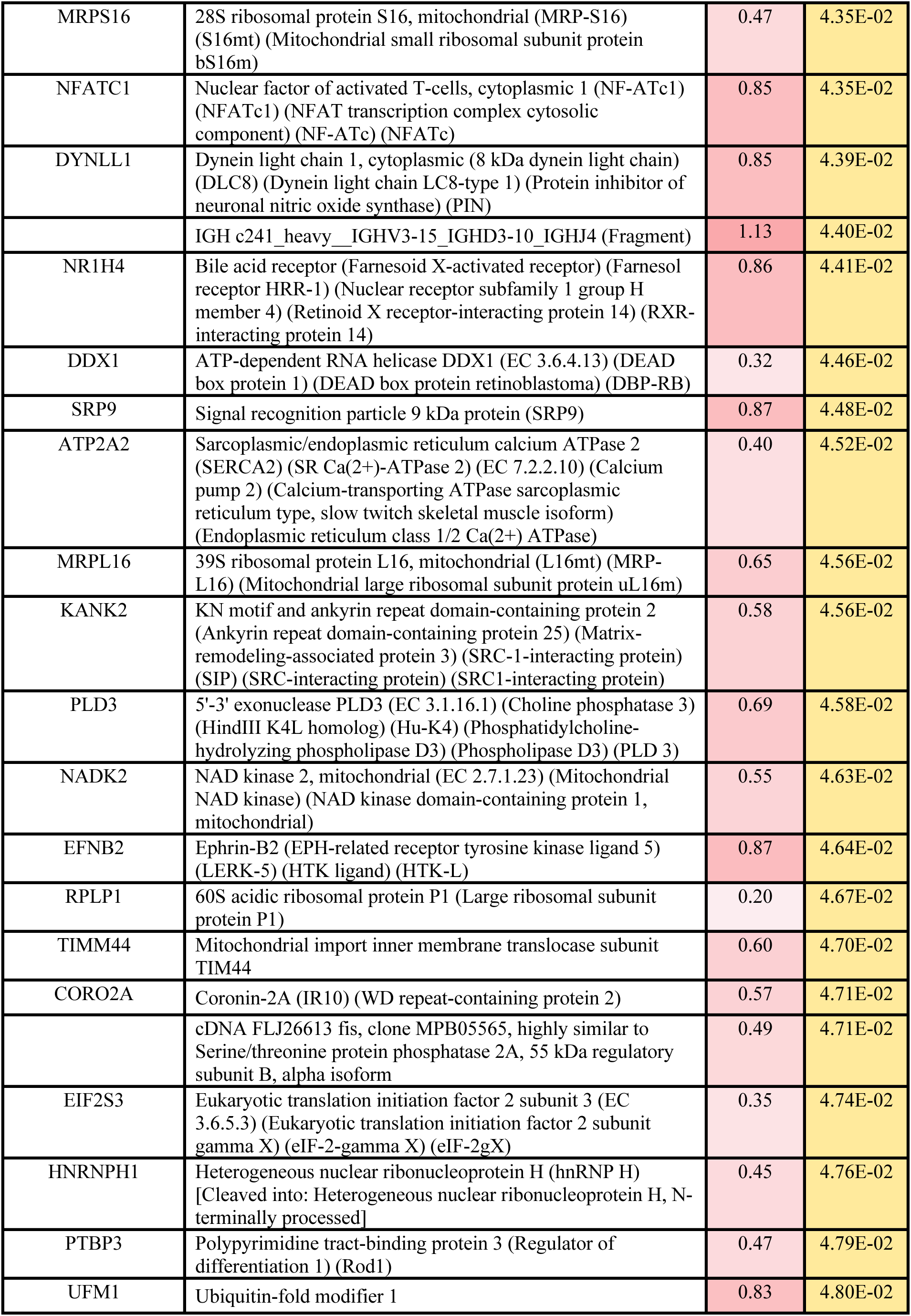
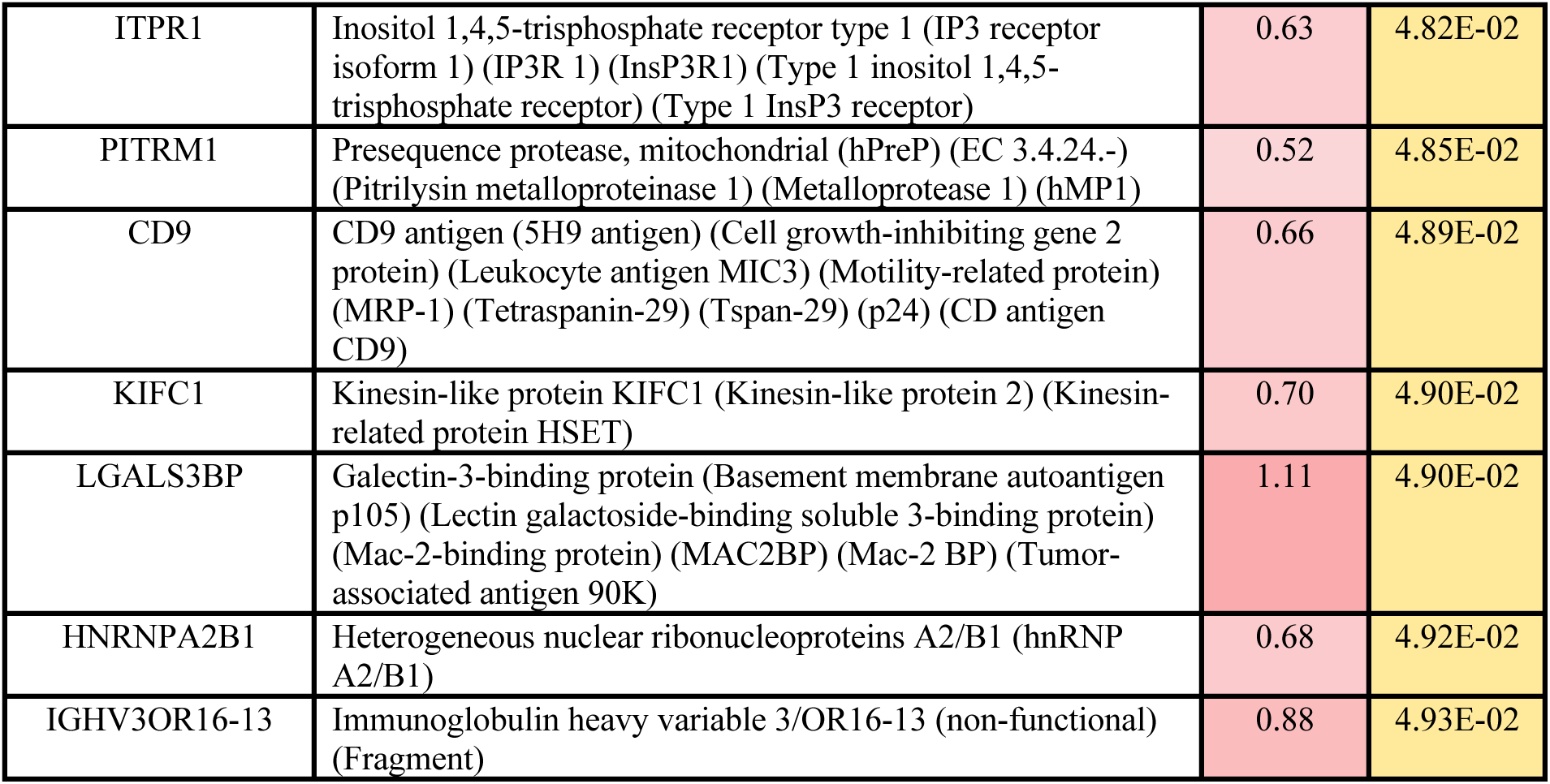
All significantly enhanced proteins associated with the trapped precursor.

**Movie S1.**

Full time-lapse movie showing the transfer of healthy mitochondria (MitoTracker Green; green) within a TNT into a cell with trapped precursor (*Su9-mScarlet-DHFR* (magenta) +100 nM MTX). Cells were imaged live using an Olympus IXplore SpinSR system. Each timeframe is a Max z-projection of 10 slices, taken every 5 minutes with 10 timepoints.

## Notes

### Competing Interest Statement

The authors have declared no competing interest.

### Summary of Updates

Minor changes and addition of authors inadvertently missed the first time around.

